# Inflammatory molecules in the glomerular endothelium in mild IgA-Nephropathy identified by single-cell and spatial transcriptome

**DOI:** 10.1101/2024.10.30.621002

**Authors:** Kumi Hasegawa, Nagako Kawashima, Ayako Kawabata, Megumi Sakakura, Naoki Onoda, Takashi Sano, Itaru Urakawa, Masahiro Matsubara, Shokichi Naito

**Author notes:** **Corresponding author: Shokichi Naito**, Department of Nephrology, Kitasato University School of Medicine, 1-15-1 Kitasato, Minami-ku, Sagamihara-shi, Kanagawa, 252-0374, Japan; **Kumi Hasegawa**, Research Core Function Laboratories, Research Division, Kyowa Kirin Co., Ltd. 3-6-6, Asahi-machi, Machida-shi, Tokyo, 194-8533, Japan.

## Abstract

Immunoglobulin A nephropathy (IgA-N) is a primary glomerulonephritis characterized by mesangial cell proliferation and expansion. It still leads to end-stage renal disease over the long course of the disease in many patients. Although glomerular endothelial cells (GE) have been implicated in the pathogenesis of IgA-N, their role remains poorly understood. We conducted single-cell and spatial transcriptomic analysis using human specimens with mild IgA-N compared with normal. We observed the significantly higher expression of several inflammation-associated molecules which have been reported and not reported to be involved in IgA-N pathogenesis, respectively. Furthermore, we show that several important molecules were also highly expressed in GE of IgA-N at protein level by tissue immunostaining. Our analysis suggests that the characteristic pathways identified in GE that involve these molecules are associated with the progression of IgA-N. These findings offer important insights on both the pathogenesis of IgA-N and its treatment at an early stage.

## Introduction

Immunoglobulin A nephropathy (IgA-N) —defined as mesangial proliferative glomerulonephritis with deposition of IgA antibodies in the renal glomerular mesangial area— is the most common primary glomerulonephritis worldwide^1,2^. The progression of IgA-N follows a process known as the “four-hit theory”, whereby immune complexes containing galactose-deficient IgA1 (Gd-IgA1) are first deposited in glomerular mesangial cells, which leads to inflammation and injury followed by glomerular damage^2^. It is thought that glomerular mesangial cells in which immune complexes containing Gd-IgA1 are deposited induce glomerular endothelial cell (GE) damage in the chronic phase of IgA-N^3^.

On the other hand, some studies suggest that damaged endothelial cells cause blood Gd-IgA1 immune complexes to permeate blood vessels and deposit in the mesangial area, resulting in the development and progression of IgA-N considering that endothelial cells are located in the innermost part of capillaries^4,5^. Additionally, studies have shown that Gd-IgA1 immune complexes enhance proinflammatory cytokines by binding to GE^2,5^. In fact, glomerular lesions are known to cause capillary damage in the acute phase of IgA-N, leading to hematuria^4^. Furthermore, in previous studies, single-cell RNA sequencing (scRNA-seq) has revealed endothelial cell damage-induced capillaritis in mice with early spontaneous IgA-N^6^. However, no reports have elucidated the role of vascular endothelial cells in the pathogenesis of human IgA-N, and the mechanism by which GE are involved in the pathogenesis or progression of the disease remains poorly understood. Thus, there is no definitive treatment for IgA-N, and the challenge of the disease progressing to end-stage renal disease over the long term in many patients still remains^7^.

ScRNA-seq, which provides transcriptomes at the single-cell level, has been used to understand cellular heterogeneity in various research studies^8,9^. While various scRNA-seq platforms are available, targeting tissues generally requires single-cell isolation in advance. This leads to loss of cellular spatial information. Unlike scRNA-seq, spatial transcriptome sequencing (ST-seq) uses tissue sections, allowing quantification of gene expression while preserving location information. Among the available ST-seq systems, 10× Genomics Visium is a convenient, widely used, commercially available ST-seq platform. However, the Visium system, which captures gene expression using circular spots, each 55 μm in diameter, contains information on several to dozens of cells per spot and is therefore not compatible with gene expression analysis of only specific cells contained in aggregates comprising several cell types, such as the renal glomerulus^10^.

Although several studies using scRNA-seq in human IgA-N have been reported, information pertaining to GE inflammation in these studies is limited^11–13^. Furthermore, no previous reports have factored spatial location information based on ST-seq in human IgA-N. In another recently published study, we integrated data from scRNA-seq and ST-seq to analyze rat kidney^14^. Applying the same approach, here we successfully defined GE clusters in scRNA-seq (Fig. 1). Use of both scRNA-seq and ST-seq enabled determination of the gene expression profile of GE, one of the cell types constituting complicated renal structures such as the glomeruli, in mild IgA-N. Furthermore, activation of characteristic inflammatory responses and new inflammation-associated molecules previously not known to be involved in IgA-N were identified in GE from the mild IgA-N specimens. Our findings suggest that the inflammatory response characteristic of IgA-N in GE is also an important event in the mechanism of IgA-N.

**Figure 1.**
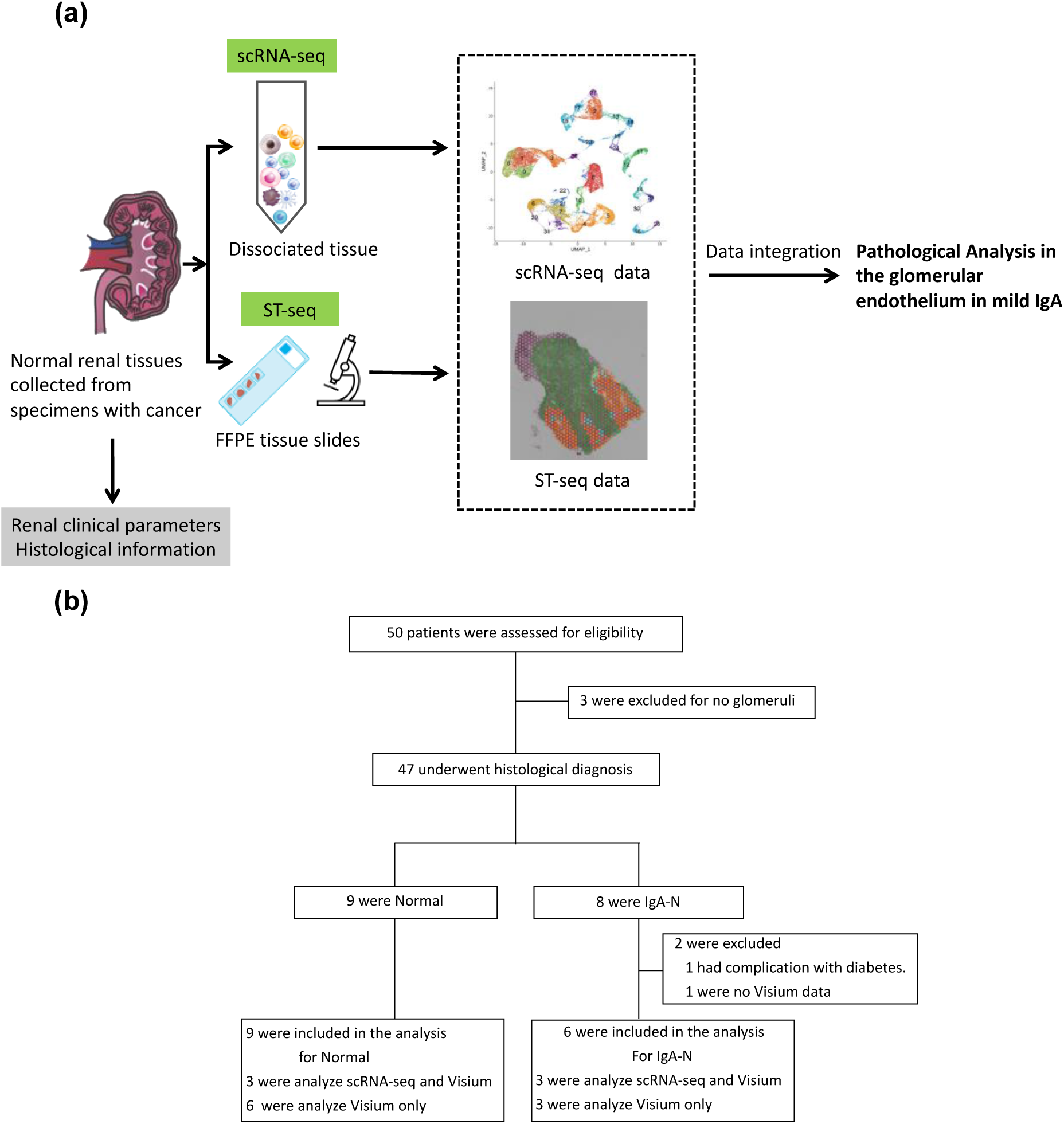
Scheme and flow diagram for integrated analysis of scRNA-seq and Visium. **(a)** Scheme and **(b)** flow diagram for integrated analysis of scRNA-seq and Visium scRNA-seq: single-cell RNA sequencing.

## Results

### Diagnosis of IgA-N

Fig. 1b depicts the flow of specimens who underwent tissue diagnosis for IgA-N in this study. The final analysis included 9 specimens with normal tissue and 6 specimens with IgA-N. All histopathological images diagnosed with IgA-N showed IgA deposition in the mesangial area and mesangial cell proliferation and mesangial matrix expansion, characteristic features of IgA-N (Fig. 2a, b, Supplementary Fig. 1a-d). Additionally, the IgA-N lesions were of grade I or II according to the Japanese histologic classification (JHC), indicating mild IgA-N^15,16^ (Table 1).

**Figure 2.**
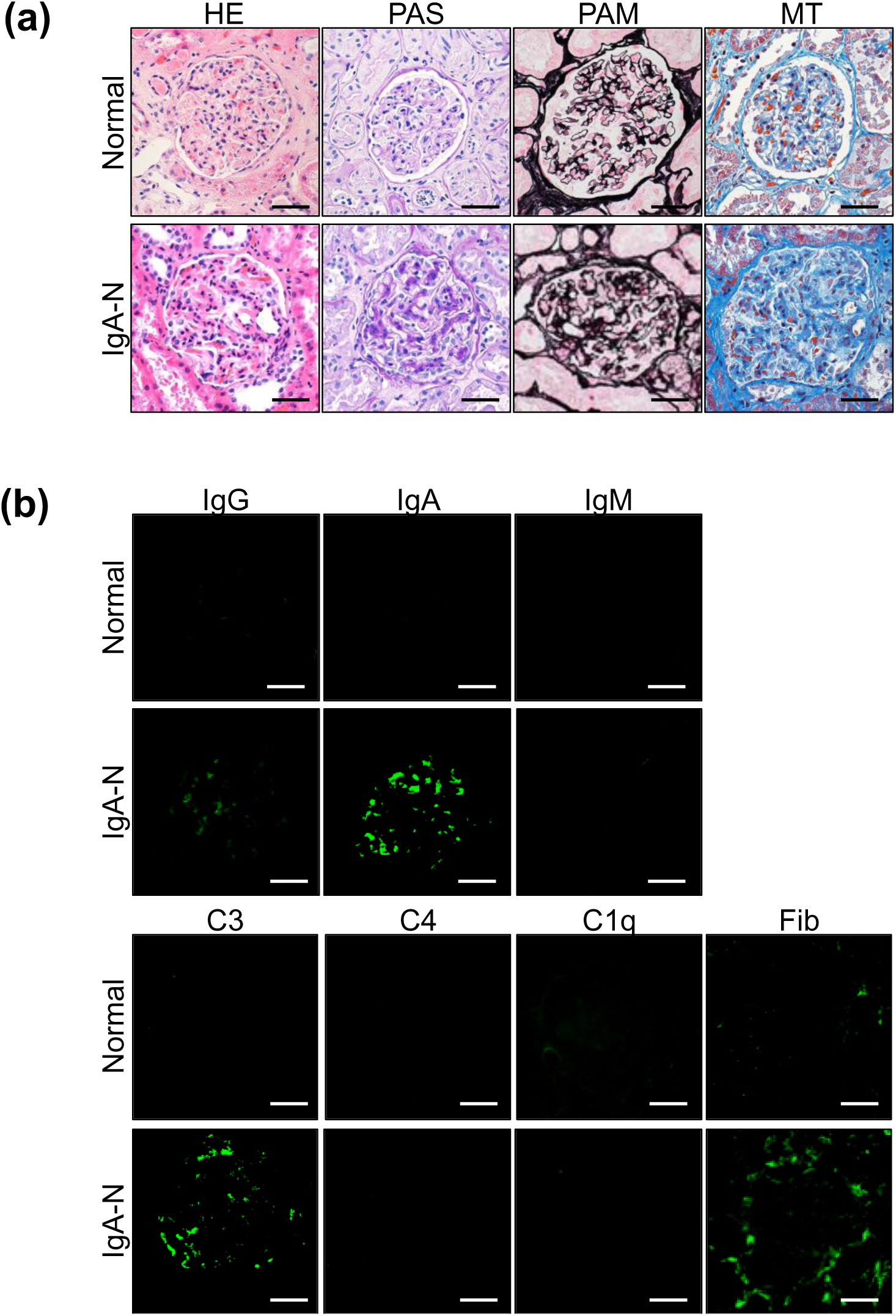
Immunostaining used for diagnosis of IgA-N. **(a)** Representative optical microscopic images of IgA-N specimens and normal glomeruli specimens, and **(b)** Immunofluorescence staining images of IgA-N specimens and normal specimens. Fib: fibrinogen; HE: hematoxylin-eosin stain; IgA: immunoglobulin A; IgA-N: IgA-Nephropathy; Masson: Masson’s trichrome stain; PAM: periodic acid-methenamine silver stain; PAS: periodic acid-Schiff stain. Scale bar = 40 µm.

**Table 1.** Baseline characteristics of samples.

|  | Normal | IgA-N | p-value |
| --- | --- | --- | --- |
| Clinical parameters |  |  |  |
| specimens, n | 9 | 6 |  |
| Male to female ratio | 6:3 | 4:2 | 0.73 |
| Age (years) | 69 (61.0, 72.5) | 65.5 (54.0, 68.0) | 0.34 |
| Height (cm) | 167.5 (156.9, 170.9) | 164.5 (160.5, 174.0) | 0.86 |
| Body weight (kg) | 61.0 (57.8, 71.4) | 72.2 (59.8, 86.8) | 0.27 |
| Serum albumin (g/dL) | 4.10 (3.75, 4.45) | 4.05 (3.90, 4.10) | 0.34 |
| Serum creatinine (mg/dL) | 0.90 (0.61, 0.98) | 0.93 (0.90, 0.97) | 0.46 |
| eGFR (mL/min/1.73 m<sup>2</sup>) | 65.0 (58.8, 73.5) | 64.0 (50.0, 67.0) | 0.53 |
| Proteinuria (g/gCr) | 0.14 (0.03, 0.21) | 0.18 (0.08, 0.32) | 0.41 |
| Serum urea nitrogen (mg/dL) | 16.5 (12.6, 20.35) | 17.7 (11.9, 20.0) | 0.95 |
| Blood glucose (mg/dL) | 115 (114.8, 130.8) | 98 (94, 100) | 0.06 |
| Hemoglobin (g/dL) | 14.7 (13.3, 14.95) | 14.45 (14.3, 15.0) | 0.95 |
| Use of antihypertensive drugs | 4/9 | 5/6 | 0.68 |
| Pathological parameters |  |  |  |
| Oxford M0/M1 lesion | 4/2 | — | — |
| Oxford E0/E1 lesion | 6/0 | — | — |
| Oxford S0/S1 lesion | 2/4 | — | — |
| Oxford C0/1/2 lesion | 1/4/1 | — | — |

|  | **Normal** | **IgA-N** | **p-value** |
| --- | --- | --- | --- |
| JHC I/II/III/IV | 5/1/0/0 | — | — |

Data are expressed as median and interquartile range (25th and 75th percentiles).
C: crescent; Cr: creatinine; E: endocapillary hypercellularity; eGFR: estimated glomerular filtration rate; IgA: immunoglobulin A; IgA-N: IgA nephropathy; JHC: Japanese histologic classification; M: mesangial hypercellularity; Normal: healthy subjects; S: segmental glomerulosclerosis.

### Cell identification using scRNA-seq

ScRNA-seq was performed using human kidney samples from 3 specimens with IgA-N and 3 normal subjects with normal tissue (Fig. 1b), yielding a total of 45,488 cells with a median of 2,375 genes per cell and a mean read count per cell of approximately 51,260 (post-normalization mean reads per cell) (Supplementary Table 2). Unsupervised clustering analysis yielded 32 cell clusters, with the following 13 cell types identified by the expression of known renal cell marker genes: GE, Cluster #18; mesangial cell (mesa), Clusters #19 and #24; podocyte (podo), Cluster #20; proximal tubule (PT), Clusters #1, #3, #8, #9, and #26; distal tubule (DT), Clusters #0, #10, and #22; distal convoluted tubule (DCT), Clusters #5 and #21; collecting duct (CD), Clusters #4, #6, #7, #25, #29, and #31; endothelial cell (Endo), Clusters #2, #13, #15, #17, and #27; smooth muscle cell (SMC), Clusters #11 and #12; B cell (B), Cluster #30; T cell (T), Cluster #16; monocyte (Mono), Clusters #14 and #23; and NK cell (NK), Cluster #28 (Fig. 3a, b, Supplementary Fig. 2, Supplementary Table 3). All clusters defined as GE were detected from both IgA-N and normal specimens (Fig. 3c).

**Figure 3.**
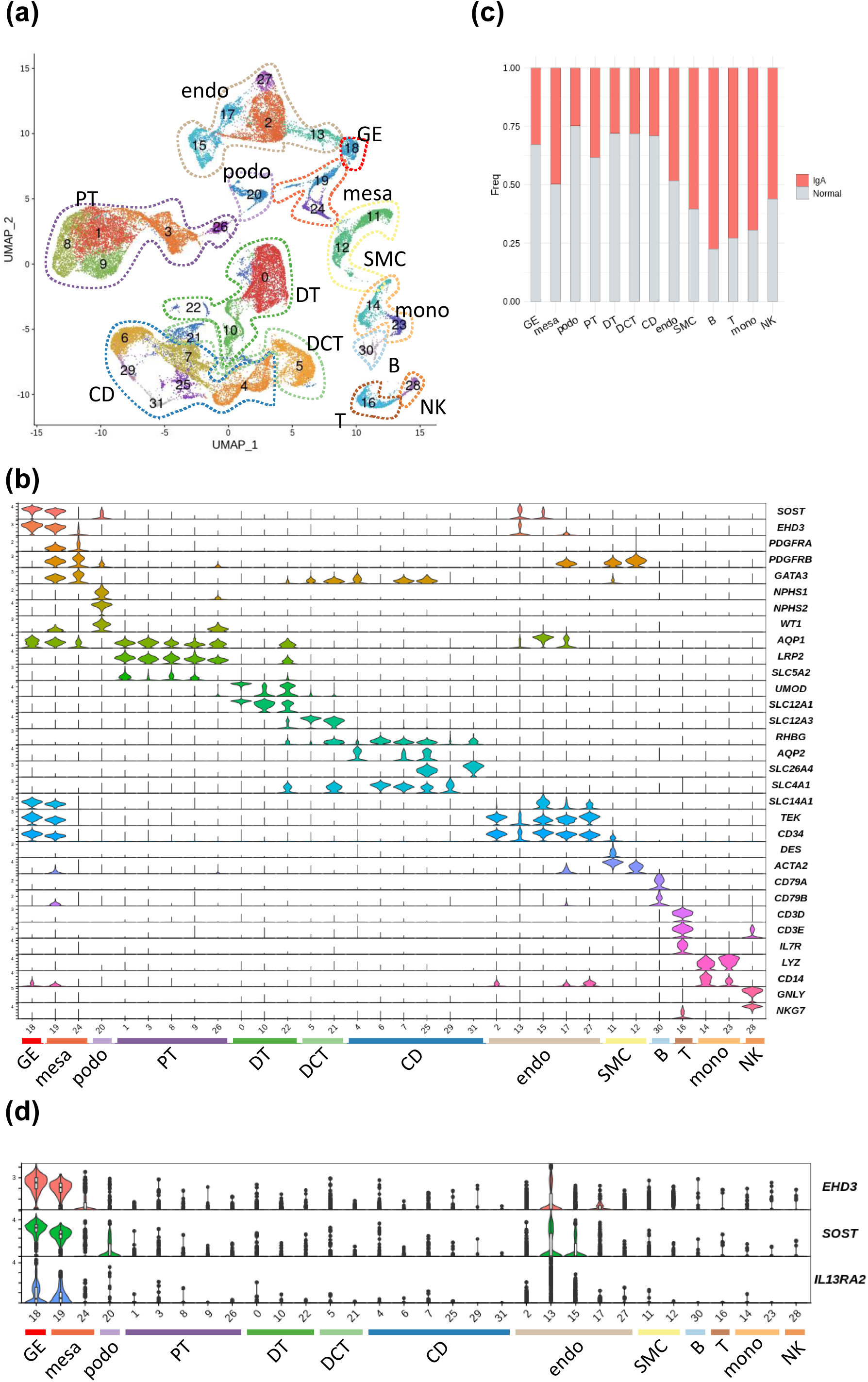
scRNA-seq of cells isolated from human renal tissues. **(a)** UMAP plots of major renal cell populations identified by unsupervised clustering and annotation with marker genes; **(b)** Violin plots showing cluster-specific expression of genes representative of each cluster of known marker genes in major renal cells. Based on the marker genes, the following cells were identified: glomerular endothelium (GE), Cluster #18; mesangial cell (mesa), Clusters #19 and #24; podocyte (podo), Cluster #20; proximal tubule (PT), Clusters #1, #3, #8, #9, and #26; distal tubule (DT), Clusters #0, #10, and #22; distal convoluted tubule (DCT), Clusters #5 and #21; collecting duct (CD), Clusters #4, #6, #7, #25, #29, and #31; endothelial cell (Endo), Clusters #2, #13, #15, #17, and #27; smooth muscle cell (SMC), Clusters #11 and #12; B cell (B), Cluster #30; T cell (T), Cluster #16; monocyte (mono), Clusters #14 and #23; and NK cell (NK), Cluster #28; **(c)** Count ratio of cells from IgA-N specimens versus cells from normal specimens for each annotated cell type; and **(d)** Violin plot showing gene expression in each cluster of *IL13RA2*, a new GE marker gene. IgA: immunoglobulin A; IgA-N: IgA nephropathy; scRNA-seq: single-cell RNA sequencing; UMPA: uniform manifold approximation and projection.

Further, both EH domain containing 3 (*EHD3*)^17^ and sclerostin (*SOST*)^18^, known GE markers, were expressed in the GE clusters of interest. The expression of interleukin 13 receptor subunit alpha 2 (*IL13RA2*), a molecule previously not known GE marker, was also high in GE clusters (Fig. 3d). Next, immunostaining of SOST and IL13RA2 using EHD3 as the index demonstrated a specific staining pattern of the GE in both IgA-N and normal specimens (Fig. 4a-d).

**Figure 4.**
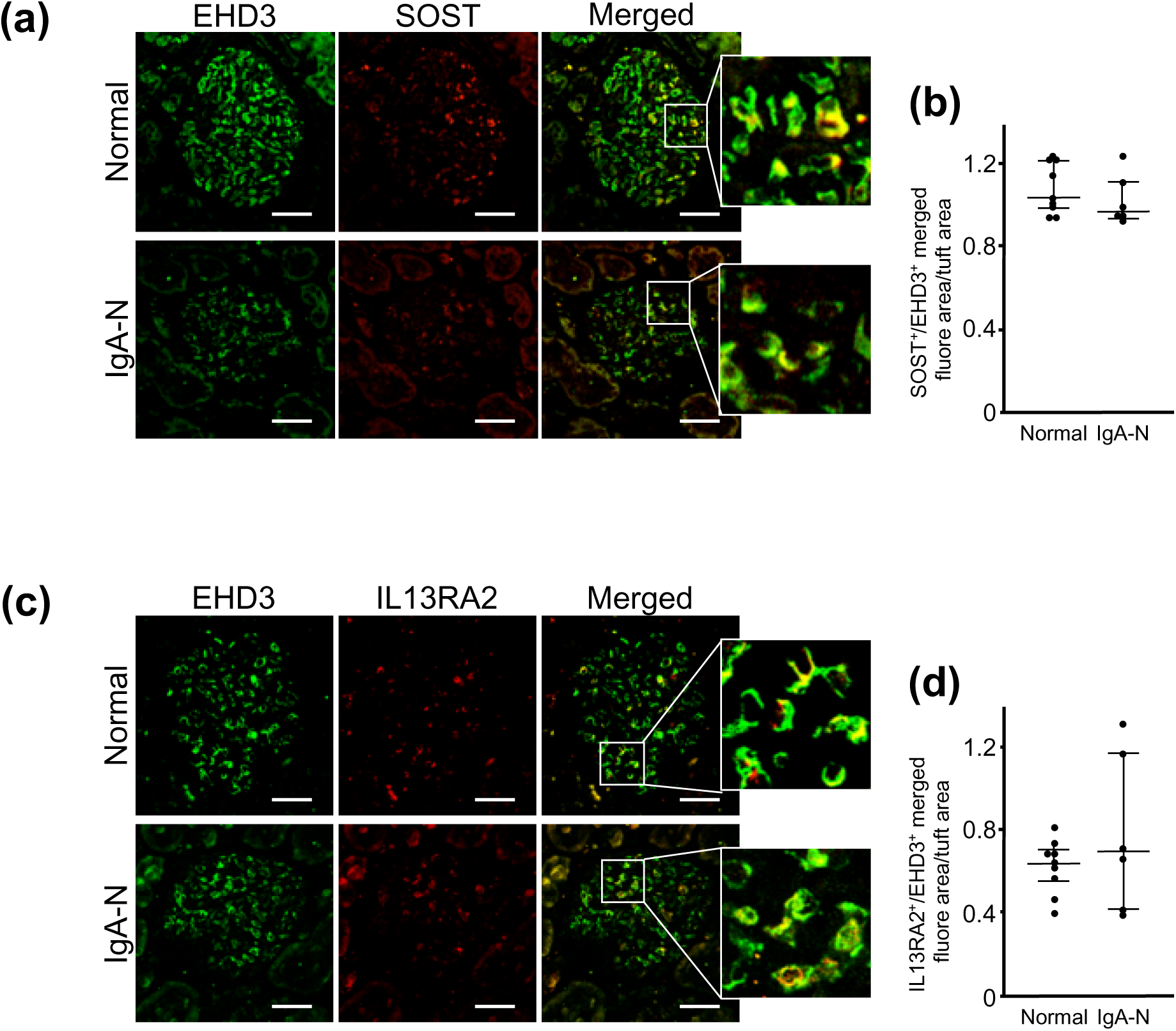
Immunofluorescence staining images of GE markers. **(a)** Double-stained images of EHD3 (green) and SOST (red), **(b)** Fluorescence intensity of SOST, **(c)** Double-stained images of EHD3 (green) and IL13RA2 (red), and **(d)** Fluorescence intensity of IL13RA2. IgA: immunoglobulin A; IgA-N: IgA nephropathy. Scale bar = 40 µm.

### Validation of spatial distribution of GE

In addition to the same specimens as used for scRNA-seq, kidney samples from 3 specimens with IgA-N and 6 normal subjects with normal tissues were used for the Visium analysis (Fig. 1b), yielding a total of 22,782 spots with a median of 4,918 genes detected from each spot (Supplementary Table 4) and a mean read count per spot of approximately 57,646 (post-normalization mean reads per spot). For spots on each section, the two renal regions of the cortex and medulla and the glomeruli were identified by experts based on pathological information (Fig. 5a, Supplementary Fig. 3, 4). Expression of representative glomerular constituent cell markers (podo; NPHS1 adhesion molecule, nephrin (*NPHS1*), mesa; platelet derived growth factor receptor alpha (*PDGFRA*), GE; *EHD3*) was confirmed in the glomerular clusters (Fig. 5c).

**Figure 5.**
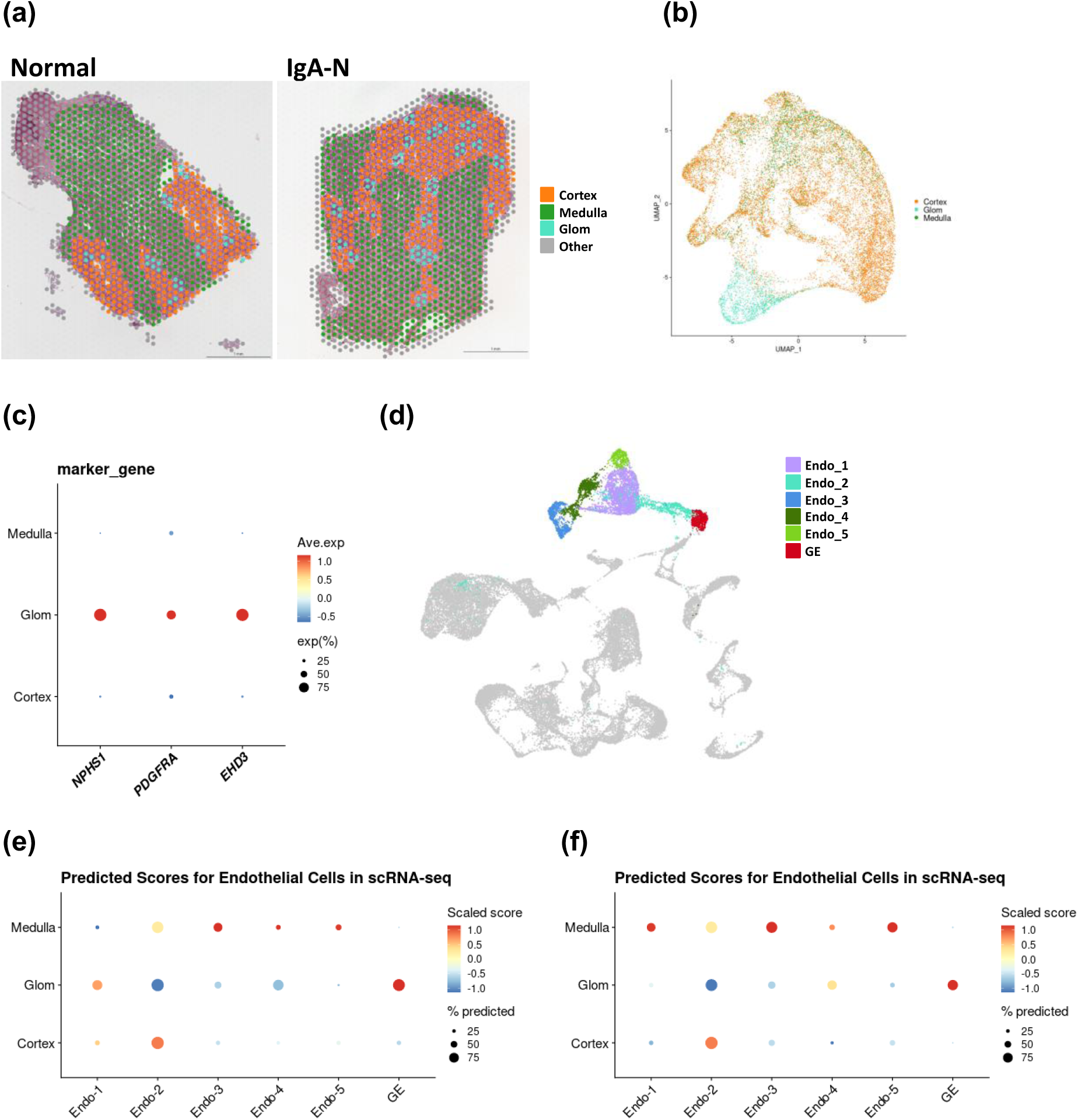
Integrated analysis of scRNA-seq and Visium. Spots detected from histological images of normal specimens and IgA-N specimens, with **(a)** UMAP plots for a pool of all samples colored with histological annotation. **(b)** Detected spots classified by a pathologist into the following four regions according to their histological features: cortex, medulla, glomeruli (Glom), and other. **(c)** Dot plots showing the expression of glomerular constituent cell marker genes (podo; *NPHS1*, mesa; *PDGFRA*, GE; *EHD3*) in each region. **(d)** UMAP plots of the clusters used for integrated analysis with Visium: GE, Cluster #18; and endothelial cells (Endo) (Endo1, Cluster #2; Endo2, Cluster #13; Endo3, Cluster #15; Endo4, Cluster #17; and Endo5, Cluster #27). Gray dots indicate cells not used for the integrated analysis with Visium. Predictive score of each cell type (scRNA-seq data; X-axis) against each histological annotation (Visium data; Y-axis) of **(e)** normal specimens and **(f)** IgA-N specimens. The dot color represents the scaled predictive score, and the dot size represents the proportion of spots with a predictive score of >0 in each histological annotation. IgA: immunoglobulin A; IgA-N: IgA nephropathy; scRNA-seq: single-cell RNA sequencing; UMPA: uniform manifold approximation and projection.

An integrated analysis of scRNA-seq and Visium was performed to confirm whether the GE population estimated in scRNA-seq was located in the glomerular area. scRNA-seq data of the endothelial cell population (Endo1, Cluster #2; Endo2, Cluster #13; Endo3, Cluster #15; Endo4, Cluster #17; and Endo5, Cluster #27) and putative GE population (Cluster #18) (Fig. 5d) were integrated with the Visium data separately for IgA-N specimens and normal specimens. The predictive scores for the GE clusters in the glomerular area were high for both IgA-N and normal specimens (Fig. 5e, f, Supplementary Fig. 5a, b).

### Differential expression analysis of the GE from mild IgA-N specimens

Functional differences in the GE between mild IgA-N specimens and normal specimens were explored.

IgA-N is characterized by proliferative changes in the mesangial cells and matrix and deposition mainly of IgA in the mesangial area, which were observed in the present specimens (Fig. 2a, b). First, we assessed whether the pathway analysis using ingenuity pathway analysis (IPA) could capture the functional differences, which observed in histopathological analysis of IgA-N specimens, between IgA-N specimens and normal specimens in the mesangial cell population. The results showed that fibrosis-associated pathways were enriched in mesangial cells from IgA-N specimens (Fig. 6a, Supplementary Table 5, 6). Histopathological images also demonstrated mesangial matrix expansion, which indicates increased mesangial fiber components in the mesangial area^19^ (Fig. 2a), a finding consistent with the results of the pathway analysis. Thus, it was confirmed that the pathway analysis in the current study was capable of capturing functional differences between IgA-N specimens and normal specimens.

**Figure 6.**
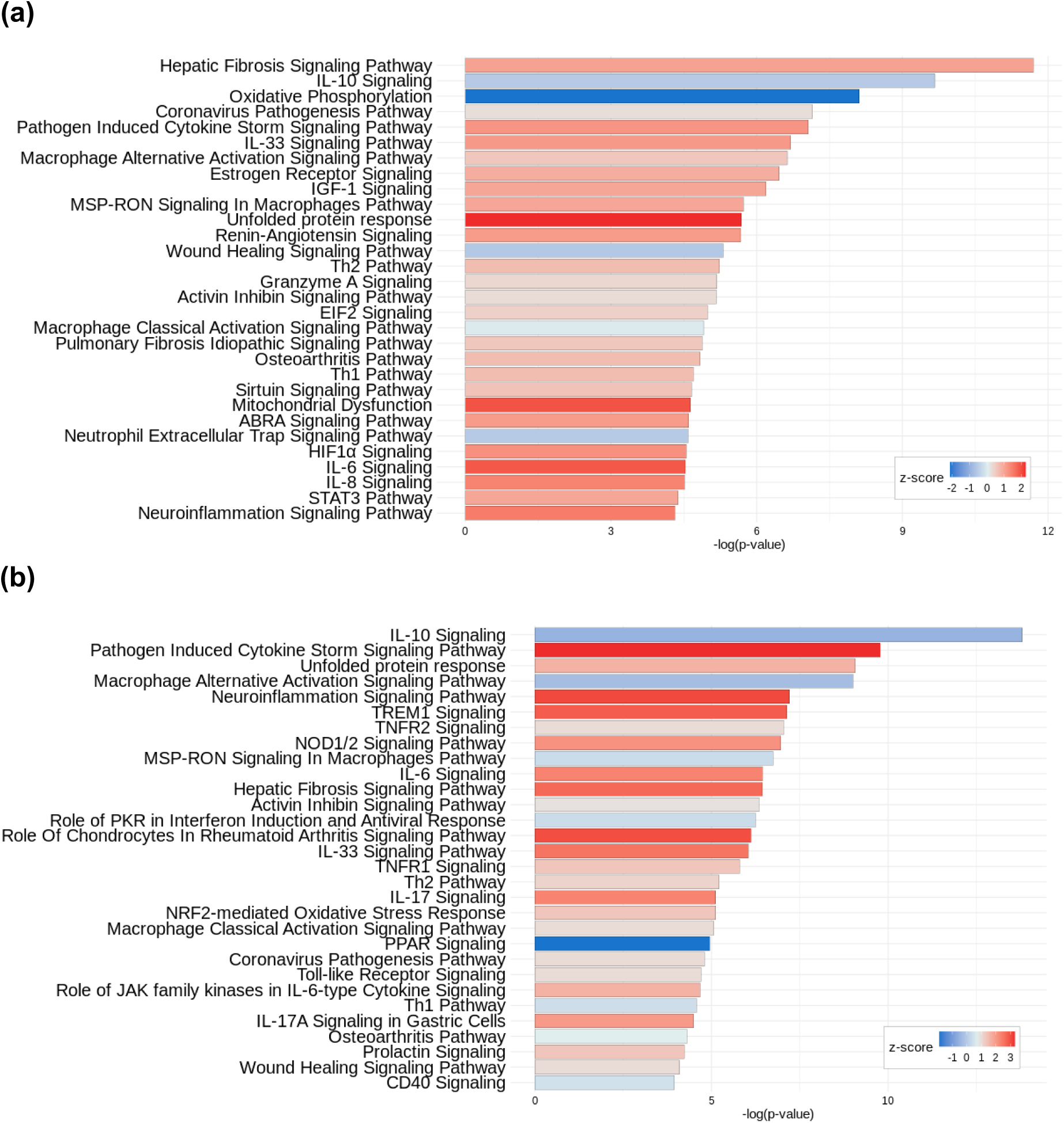
Pathway analysis of mesangial cells and GE. A pathway list was prepared for **(a)** mesangial cells and **(b)** GE using DEGs with a log-fold change of ≥0.25 in IgA-N specimens compared with normal specimens. The 30 pathways with -log_­­­_(p-value) are listed in descending order. The pathways with no z-score determined were excluded. DEG: differentially expressed gene; IgA: immunoglobulin A; IgA-N: IgA nephropathy.

A similar pathway analysis was performed using GE from IgA-N specimens and normal specimens. Inflammation-associated pathways such as interleukin-6 (IL-6) signaling were found to be enriched in GE from IgA-N specimens (Fig. 6b, Supplementary Table 5, 6). The differentially expressed genes with a z-score of >2 contained in the inflammation-associated pathways included representative inflammation-associated molecules such as C-C motif chemokine ligand 2 (*CCL2*) and C-X-C motif chemokine ligand 2 (*CXCL2*) (Supplementary Table 5). Additionally, immunohistochemistry (IHC) of CCL2 and CXCL2 was performed to confirm whether GE from the IgA-N specimens were inflamed. CCL2 and CXCL2 expression was found to be elevated in the GE from IgA-N specimens (Fig. 7a-d). scRNA-seq and IHC yielded consistent results. Among the differentially expressed genes with a z-score >2 contained in the inflammation-associated pathways, increased expression of endothelin receptor type B (*EDNRB*), Interleukin 1 receptor-like 1 (*IL1RL1*), phospholipase C gamma 2 (*PLCG2*), and CCAAT enhancer binding protein beta (*CEBPB*), previously not known to be associated with human IgA-N, was noted (Supplementary Table 7). Visium data showed no increase in the expression of any of these genes (Supplementary Table 8).

**Figure 7.**
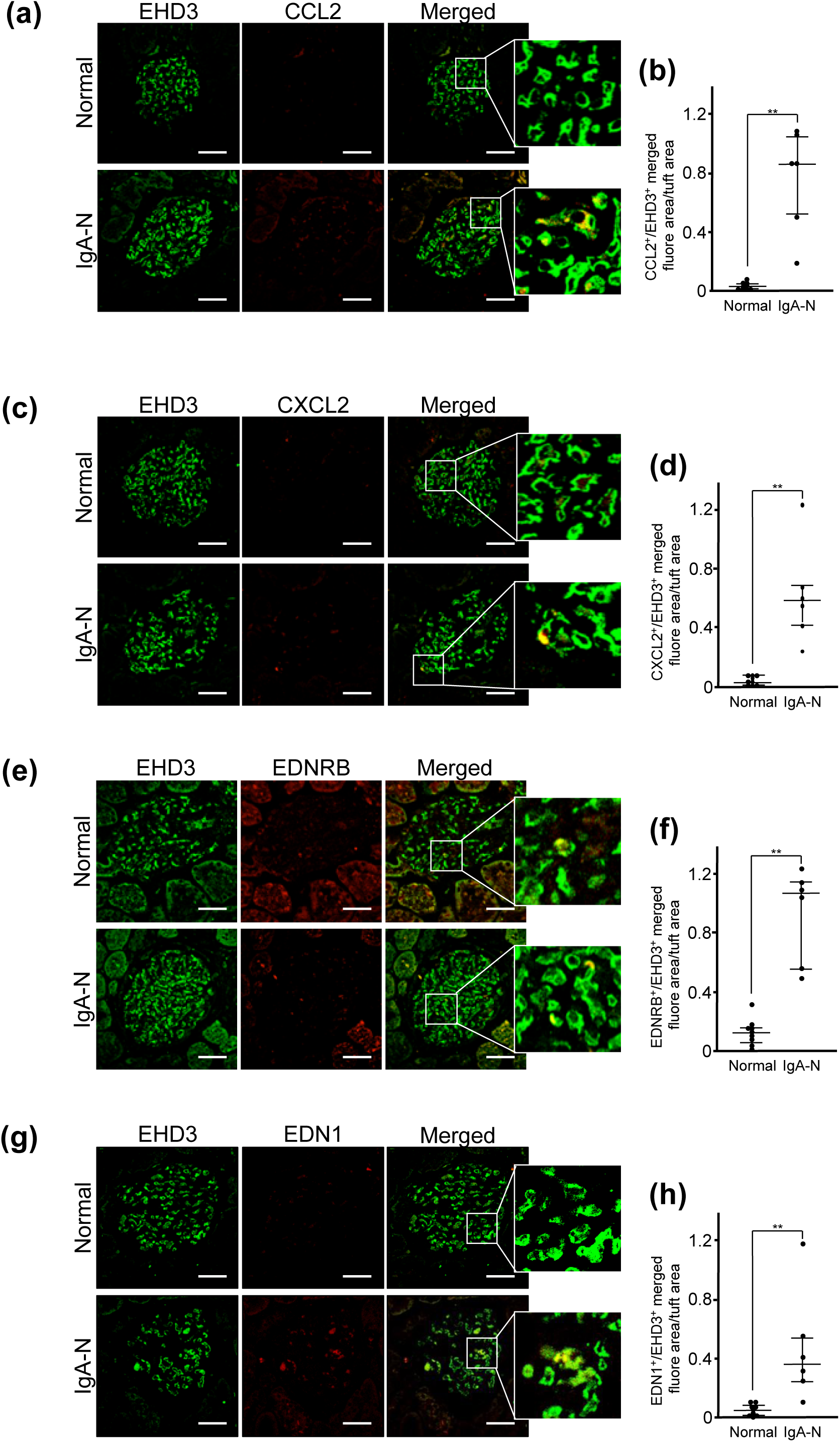
Immunofluorescence staining images and fluorescence intensity in the glomeruli of inflammation-associated factors in normal specimens and IgA-N specimens. **(a)** Double-stained images of EHD3 (green) and CCL2 (red); **(b)** Fluorescence intensity of CCL2; **(c)** Double-stained images of EHD3 (green) and CXCL2 (red); **(d)** Fluorescence intensity of CXCL2; **(e)** Double-stained images of EHD3 (green) and EDNRB (red); **(f)** Fluorescence intensity of EDNRB; **(g)** Double-stained images of EHD3 (green) and EDN1 (red); and **(h)** Fluorescence intensity of EDNRB. IgA: immunoglobulin A; IgA-N: IgA nephropathy. Scale bar = 40 µm; **p < 0.01.

*EDNRB*, the most significantly differentially expressed gene, was examined for protein expression level and localization by immunostaining. Consistent with the scRNA-seq results, increased expression of EDNRB was confirmed in the GE from IgA-N specimens (Fig. 7e-f). Similarly, expression of endothelin 1 (EDN1), the ligand for EDNRB, was also significantly increased in the GE from IgA-N specimens (Fig. 7g, h).

## Discussion

In this study, we identified the gene expression profile of GE using scRNA-seq and Visium in renal tissues from human specimens with mild IgA-N compared with normal specimens. Analysis of the cellular status in mild IgA-N may be informative for early therapeutic interventions, in that the pathogenesis and degree of progression becomes known at an early stage of the disease. We performed an integrated analysis of scRNA-seq and Visium to clearly define GE based on scRNA-seq data. Comparison of the gene expression profiles in GE between mild IgA-N specimens and normal specimens demonstrated that inflammation-associated pathways were activated in GE in mild IgA-N. Additionally, several inflammation-associated factors previously not known to be involved in IgA-N were identified. Thus, this study provides new insights into the mechanisms of GE in mild IgA-N in humans.

IgA-N is a prevalent disease in the Asian region^2^ and is estimated to develop in 0.04% of the Japanese population^20^. The current analysis using specimens from Japanese patients with renal cell carcinoma showed that as high as approximately 17% of the patients had IgA-N (Fig. 1b). Previous studies have reported an incidence of 2.9%-18.3% based on glomerular analysis of isolated kidney samples^21,22^ and have also suggested the possibility of some forms of IgA-N being associated with renal cell carcinoma^21^. Although the incidence of IgA-N in association with renal cell carcinoma may have been high in the current study as well, the pathogenesis of IgA-N is considered to have followed the “four-hit theory”, in which a similar process develops as a result of various causes^2^. Furthermore, considering that the mean age at diagnosis of renal cancer in the general population is 65 years^23^, similar to that observed in the current analysis population, the time from onset of IgA-N in our study was much shorter than that observed in cases of mild IgA-N that occur at a young age and slowly progress^23^. Thus, the possibility that endothelial cell inflammation, described later, occurs early in IgA-N cannot be ruled out.

We attempted to understand the role of GE in the pathogenesis of IgA-N based on scRNA-seq data. The integrated analysis of scRNA-seq and ST-seq indicated that the GE clusters identified by the expression of the marker genes in scRNA-seq were predicted to localize at glomeruli. The IHC staining results of molecular markers using the same samples also showed expression of these markers in GE, establishing the cell group as the GE. EHD3 and SOST were also identified as novel marker molecules of GE in mild IgA-N.

Furthermore, *IL13RA2* was newly identified as a gene specifically expressed in GE. *IL13RA2* has previously not been described in association with glomerular disease. Of note, IL13RA2 was expressed in a subset of GE, regardless of the presence or absence of IgA-N (Fig. 4c, d). Several publicly available normal human renal scRNA**-**seq datasets containing GE have also shown expression of *IL13RA2* in GE, regardless of age (Supplementary Fig6a-e.), suggesting that *IL13RA2* may serve as a GE marker in normal specimens.

Additionally, we showed that inflammatory responses were activated in the GE in mild IgA-N. Although previous studies have reported activated inflammatory responses in the GE in mild IgA-N, the findings pertained to only specific molecular species^5,24,25^, and to the best of our knowledge, this is the first study to demonstrate activation of GE inflammatory responses based on expression analysis at single-cell resolution with spatial context.

IgA-N often presents with hematuria after upper respiratory tract infection, and its pathology is believed to involve inflammation associated with bacterial infection^2^. The current study also showed activation of the signal pathways associated with proinflammatory cytokines such as IL-6 (Fig. 6). Gd-IgA1 immune complexes promote migration of inflammatory cells by increasing adhesive factors such as intracellular adhesion molecule 1 (ICAM-1) and induce the production of proinflammatory cytokines by GE, thereby exacerbating intrarenal inflammation^5^. Moreover, both GE and mesangial cells associated with Gd-IgA1 immune complexes secrete IL-6 that induces loss of endothelial cells, increases endothelial permeability, and recruits inflammatory cells, resulting in enhancement of Gd-IgA1 deposition and inflammatory responses in the mesangial cell^19,25^. Several genes other than ICAM-1 that are known to be associated with mild IgA-N, including selectin E (*SELE*)^26^ and C-X3-C motif chemokine ligand 1 (*CX3CL1*)^6^, have shown increased expression in GE in a mouse model of mild IgA-N (gddY mice). These findings suggest that endothelial cell inflammation may also be involved in the mechanism of pathogenesis and progression of human IgA-N, consistent with the results in the mouse model of IgA-N^6^. Furthermore, given that the gene expression profiles of GE from gddY mice and those from humans with mild IgA-N appear to be partly similar, gddY mice may be useful for functional analysis of the newly identified genes and proteins.

We also identified *EDNRB*, *CEBPB*, *IL1RL1*, and *PLCG2* as inflammation-associated genes previously not known to be associated with human IgA-N.

EDN1 is a bioactive peptide with a blood flow-regulating effect and has two types of receptors, type A and type B^27^. Endothelin receptor type A (EDNRA), located on vascular smooth muscle cells, is one of the most potent vasopressor substances in nature^27^, and Sparsentan, a dual antagonist of EDNRA and angiotensin receptors, has a therapeutic effect on IgA-N^28^. On the other hand, EDNRB, expressed in vascular endothelial cells, exerts vasodilatory effects^29^. The current scRNA-seq analysis showed no difference in the expression level of either *EDN1* or *EDNRA* between IgA-N and normal tissues, but there was an increased expression of *EDNRB* in specimens from specimens with IgA-N (Supplementary Table 5). Tissue staining revealed elevated expression of EDN1 in the glomeruli from patients with IgA-N, as previously reported^30^. A possible reason is not the result of increased mRNA expression in GE, but rather reflects the result of serum EDN1 binding to EDNRB. In addition, the mild IgA-N cases in our study might have had increased expression of *EDNRB*, which has vascular protective effects, as a compensatory mechanism for glomerular damage.

Our IgA-N tissue specimens also showed increased expression of *PLCG2* (Supplementary Table 5). Although the role of PLCG2 in IgA-N remains unknown, it has been reported that activation of PLCG2 activates the 1,2-diacylglycerol-protein kinase C (PKC) pathway^31,30^, which regulates vascular permeability^32^. Thus, it is also possible that enhancement in vascular permeability by PLCG2 may induce Gd-IgA1 deposition in the mesangial area.

Further, we observed elevated expression of the chemokine-related genes *CCL2* and *CXCL2* in the IgA-N specimens (Supplementary Table 5). Although CCL2 and CXCL2 are known to be associated with IgA-N^33,34^, elevated expression of these molecules in the GE in IgA-N has not been described previously. Chemokines in endothelial cells are involved in various kidney diseases as inflammatory mediators^35^. CCL2, also known as monocyte chemoattractant protein-1 (MCP-1), is a member of the chemokine family, and its expression is elevated in the glomeruli in IgA-N^36,37^. Hypertension and diabetes also increase CCL2 expression in renal endothelial cells^37,38^. However, considering that the study excluded patients with diabetes and that similar numbers of subjects had received antihypertensive drugs in the normal group and IgA-N group, the increase in CCL2 expression could be attributable to IgA-N. Although *CXCL2* is elevated in renal tissue from patients with IgA-N^39^, its expression status in GE is unknown. One study reported that *CXCL2* expression is also increased in human aortic endothelial cells (HAECs) stimulated by interleukin 17A (IL-17), which is implicated in various inflammatory diseases^40^. Similar to this report, the current analysis revealed activation of the IL-17-related signaling pathway (Fig. 6b), suggesting that *CXCL2* may also be involved in the progression of IgA-N.

In diabetic nephropathy, a disease in which the role of GE dysfunction has attracted attention, platelet derived growth factor receptor beta (*PDGFR*) expression has been shown to be elevated in GE^41^. However, the current differentially expressed gene analysis detected no activation of either the vascular endothelial growth factor (*VEGF*) or platelet derived growth factor (*PDGF*) signaling pathway. Furthermore, although proinflammatory cytokines (interleukin 6 (IL-6) and interleukin 8 (IL-8)) are released in anti-MPO-ANCA antibody- or anti-PR3-ANCA antibody-associated crescentic glomerulonephritis, a representative glomerular disease caused by endothelial cell damage^42^, the current study found an increase in only IL-8. These data suggest that the signaling pathways in endothelial cell damage caused by IgA-N are distinct from those in specimens with diabetes or crescentic glomerulonephritis.

Thus, integration of scRNA-seq and Visium analyses revealed a tendency of more active inflammatory responses in GE than in mesangial cells in glomerular cells from specimens with mild IgA-N (Supplementary Table 9). In gddY mice, GE have been reported to play a critical role in the early phase of intraglomerular inflammation in IgA-N^6^. In fact, treatment for IgA-N generally involves steroids with anti-inflammatory effects^2^. Thus, the inflammation-associated signaling pathways in GE characteristic of IgA-N identified in the current study are not only valuable for understanding the progression of IgA-N but also offer insights into the future development of therapeutic agents.

Some limitations of the study need to be acknowledged. First, the absolute number of specimens available for analysis was small, particularly in the IgA-N group. As mild IgA-N lacks prominent symptoms such as proteinuria, subject recruitment depends on chance. To overcome the heterogeneity and individual differences in the pathology of IgA-N, large-scale recruitment is needed in future studies to ensure a larger sample size of specimens with mild IgA-N. Additionally, the study used organs resected from specimens with cancer after removal of cancer sites, which may still have had influences of cancer tissue. While the study successfully captured genes present in IgA-N by analyzing the differences between IgA-N specimens and normal specimens, determination of whether or not this gene group contributes to IgA-N needs further investigation, including *in vivo* and *in vitro* experiments and studies using tissues from specimens other than those with cancer. Second, some differentially expressed genes captured by scRNA-seq were not captured by Visium-based analysis. A possible reason is that the granularity of transcriptional profile data differs between Visium and scRNA-seq. As Visium spots contain multiple non-GE, differential expression in GE may have been obscured. Other recently available high-resolution spatial transcriptome technologies such as Xenium, VisiumHD, and CosMx may be useful for analysis of cells constituting complicated structures such as the glomeruli.

Nevertheless, the study demonstrated that GE damage is key even in patients with mild IgA-N, providing invaluable data for future studies on IgA-N. Moreover, we established a role of the inflammation-associated molecules in GE in mild IgA-N, providing new information not only for elucidating the mechanisms of IgA-N pathogenesis but also for developing therapeutic strategies for suppression of inflammation.

## Methods

### Sample acquisition and diagnostics

All renal tissues were obtained from specimens who had undergone partial nephrectomy or radical nephrectomy for renal cancer at Kitasato University Hospital (Department of Urology) and had provided written informed consent.

### Antibodies

Anti-IgG, IgA, IgM, C3, C4, C1q, fibrinogen, EHD3, EDN1, SOST, IL13RA2, CCL2, CXCL2, EDNRB, and fluorescence-labeled secondary antibody corresponding to the primary antibody were used to diagnose IgA-N and normal specimens and determine the expression levels of each antigen. Supplementary Table 1 provides detailed information on the antibodies used.

### Tissue staining

Paraffin sections were subjected to periodic acid-Schiff’s staining, hematoxylin & eosin staining, periodic acid-methenamine silver staining (Jones’ stain), and Masson’s trichrome staining^43,44^. Immunofluorescence staining for each antigen was performed on frozen sections. Renal tissues were embedded in Tissue Tek OCT compound (Sakura Finetek Japan Co., Ltd., Tokyo, Japan) and sliced into 4-μm-thick sections. The sections were washed, incubated with the primary antibody, and stained with the secondary antibody. Pathologic images were visualized using an optical microscope (BX51; Olympus Corporation, Tokyo, Japan) and analyzed using ImageJ software (https://imagej.nih.gov/ij/). Fluorescence images were visualized using a confocal laser microscope (LSM710; Carl Zeiss AG, Oberkochen, Germany) and analyzed using ZEN imaging software (Carl Zeiss AG, Oberkochen, Germany).

### Assessment of glomerular lesions

Renal tissues were evaluated by two nephrologists. Histologic lesions of IgA-N were classified according to the 2016 Oxford and JHC classifications^45,46^. Tissues showing no immunoglobulin or complement deposition in the glomeruli or morphological changes were considered normal and used for the analysis.

### Preparation of a library for Visium analysis

For Visium analysis, a library was constructed using resected renal tissues from 32 adult specimens with cancer after removal of the cancerous part. The sequencing library was prepared using Visium Spatial for FFPE Gene Expression Kit, Human Transcriptome (10x Genomics, Inc., California, U.S.) according to the manufacturer’s instructions (CG000407_Rev_C). A histological image of each sample was acquired using an inverted microscope (Nikon Eclipse Ti; Nikon Solutions Co., Ltd. Tokyo, Japan). The library was sequenced using Illumina NextSeq2000 (Illumina, Inc. California, U.S.).

### Tissue dissociation for scRNA-seq

For scRNA-seq, renal tissues were collected from 14 adult specimens with cancer after removal of the cancerous part. Renal tissue was sectioned, dissociated in Dulbecco’s modified Eagle medium (Nacalai Tesque, Inc., Kyoto, Japan) containing 100 U/mL Collagenase D (F. Hoffmann-La Roche Ltd; [1108886600]) and 5 U of DNase I (Takara Bio Inc., Shiga, Japan) by shaking at 37°C for 60 minutes, and then filtered through a 40-μm cell strainer (Corning Inc., New York, U.S.). The resultant cell suspension was subjected to density-gradient centrifugation using Optiprep (Abbott Diagnostics Technologies AS, Oslo, Norway) and 1% (w/v) bovine serum albumin containing phosphate-buffered saline (Nacalai Tesque, Inc., Kyoto, Japan) to isolate cells and then filtered through a 35-μm cell strainer (Corning Inc., New York, U.S.).

### Preparation of a library for scRNA-seq

For each sample, the cell mass and viability were determined, and microscopy was used to confirm that there were no aggregated cells or debris. An scRNA-seq library was prepared using Chromium Next GEM Automated Single Cell 3′ Reagent Kits v3.1 (10x Genomics, Inc., California, U.S) according to the manufacturer’s instructions (CG000286 Rev_E). The library was sequenced using Illumina NextSeq2000(Illumina, Inc. California, U.S.).

### Data processing of the Visium dataset

Sequence results (BCL files) were converted to FASTQ files and mapped to the human reference genome (GRCh38) using Space Ranger (v.1.3.1; 10× Genomics; https://www.10xgenomics.com/support/software/space-ranger/downloads/previous-versions). Downstream analysis was performed using R package, Seurat (v.3.2.2; https://satijalab.org/seurat/articles/install.html). Outputs from Space Ranger were converted to Seurat objects and preprocessed using the SCTransform^47^ function. The objects of all samples were integrated using the Seurat anchor-based integration workflow^48^, followed by principal component analysis, dimensional reduction, and unsupervised clustering.

### Data processing of the scRNA-seq dataset

Sequence results (BCL files) were converted to FASTQ files and mapped to the human reference genome (GRCh38) using Cell Ranger (v.4.0.0; 10× Genomics; https://www.10xgenomics.com/support/software/cell-ranger/downloads/previous-versions). Downstream analysis was performed using R package, Seurat (v.3.2.2). The output matrices from Cell Ranger were transformed into Seurat objects, and the barcodes with <200 genes were excluded as part of primary quality control to exclude low-quality cells or cell-free droplets. After primary quality control, each object was normalized and scaled according to the Seurat standard workflow, followed by principal component analysis, uniform manifold approximation and projection (UMAP), dimensional reduction, and unsupervised clustering, based on the top 2,000 highly differentially expressed genes. Among the resultant clusters, those with a high proportion (%) of mitochondrial genes (median of mt >85%) and a low gene number (median of nFeature <1,500) were excluded, and dead cells were removed as part of secondary quality control. After secondary quality control, all objects were reprocessed according to the Seurat standard workflow and subsequently integrated by the Seurat anchor-based integration workflow, followed by principal component analysis, dimensional reduction, and unsupervised clustering (ndim34; granularity, 0.8). Cell type was identified on the basis of unsupervised clustering and known marker gene expression. Differentially expressed genes in the IgA-N group as compared with the normal group were identified in FindAllMarkers with the default parameter.

### Integration of Visium and scRNA-seq datasets

Integration of Visium and scRNA-seq datasets was performed according to Seurat’s anchor-based integration workflow^48^. Among the scRNA-seq datasets, only those of GE and endothelial cell clusters were used as reference for integration. The integrated analysis was performed using Visium and scRNA-seq datasets of the same specimens. Predictive scores for each cell type were calculated for all spots in the Visium dataset.

### Gene ontology enrichment analysis

Canonical pathway analysis was performed using QIAGEN IPA (QIAGEN Inc., https://digitalinsights.qiagen.com/IPA)^49^.

### Expression analysis of IL13RA2 using publicly available data

The figure/analysis was created with BBrowserX/Talk2Data software (Bio Turing Inc., San Diego, CA, USA). The following data, registered in BBrowserX/Talk2Data, were used for analysis: ID:LAKEETAL^50^; GSE140989^17^; ID:PMID31604275_Mature^51^; GSE160048^52^; and ID:GSE159115_6_BENIGN^18^.

### Statistics and reproducibility

Baseline values were expressed as medians (25th percentile to 75th percentile). Sex ratio was compared using the chi-square test, and for other baseline values and expression levels determined by immunostaining, group medians were compared using the Mann-Whitney U test. Statistical analysis was performed used StatFlex (v.7; Artech Co., Ltd., Tokyo, Japan).

### Approval for research

Renal tissues were obtained from specimens who had undergone nephrectomy following written informed consent under the approval of the Kitasato University Ethics Committee (KMEO) (#KMEO B20-263) and Kyowa Kirin Co., Ltd. Research Ethics Committee (# 2020_033). Subjects were identified by number, not by name, and all data were anonymized. The study was conducted in compliance with the principles of the Declaration of Helsinki.

### Data availability

All sequencing data supporting the results and Histology results of Visium in the study are deposited in the Japanese Genotype-phenotype Archive (JGA) with accession number JGAS000736.

## Acknowledgements

The authors thank all the subjects who participated in this study. The authors also thank Dr. Ishii D., Dr. Ikeda M., and Dr. Kitajima K. for help with the nephrectomy procedure; Ms. Ishigaki N. for help with tissue staining; Dr. Murakami T. for support in establishing the experimental system of sample dissociation for scRNA-seq; Mr. Miyazawa T. for managing the server for the analysis; and Dr. Yoneya T. for data quality check. Editorial assistance was provided by Cactus Life Sciences (a part of Cactus Communications) and funded by Kyowa Kirin Co., Ltd.

## Author contributions

K. H., N. K., A. K., M. S., I. U., M. M., and S. N. contributed to the conception and design of the study. N. K. and S. N. performed sample preparation for scRNA-seq and Visium analysis. N. K. captured the histopathology images, and T. S. and S. N. performed pathological assessment of the tissues. S. N. provided specimen samples and clinical data. K. H., A. K., and M. S. performed scRNA-seq and Visium experiments. K. H., A. K., and M. S. performed the data processing, database construction, and data analysis. K. H., N. K., A. K., N. O., M. M., and S. N. drafted the manuscript.

