## Supplemental Figure and Table for "Inflammatory molecules in the glomerular endothelium in mild IgA-Nephropathy identified by single-cell and spatial transcriptome": SupplementaryInformation_IgANseq.pdf

***Corresponding author**

### This file includes the following:

#### Supplementary Figures

**Supplementary Figure 1: Histological staining images**

**Supplementary Figure 2: scRNA-seq data of cells isolated from each human renal tissue specimen. UMAP plots of major renal cell populations identified by unsupervised clustering and annotation with marker genes**

**Supplementary Figure 3: Visium HE images of each human renal tissue specimen**

**Supplementary Figure 4: Visium data of spots on tissue from each human renal tissue specimen**

**Supplementary Figure 5: Integrated analysis of scRNA-seq and Visium.**

**Supplementary Figure 6: Confirmation of IL13RA2 expression in the publicly available data of human renal tissue scRNA-seq.**

#### Supplementary Tables

**Supplementary Table 1: Details of antibodies used in the study**

*The other supplemental tables (2-9) below are supplied by another file in csv format.

**Supplementary Table 2: General statistics of scRNA-seq data**

**Supplementary Table 3: Cluster-specific genes (scRNA-seq data)**

**Supplementary Table 4: General statistics of ST-seq data**

**Supplementary Table 5: DEGs for normal status versus pathological status for each cell**

**type (scRNA-seq data)**

**Supplementary Table 6: Pathways in mesangial and GE**

**Supplementary Table 7: List of differentially expressed genes with a z-score of >2 contained in the inflammation-associated pathways in glomerular endothelial cells from IgA nephropathy specimens**

**Supplementary Table 8: DEGs in glomerular spots in IgA-N (ST-seq data)**

**Supplementary Table 9: Comparison analysis of mesangial and GE**

(a)

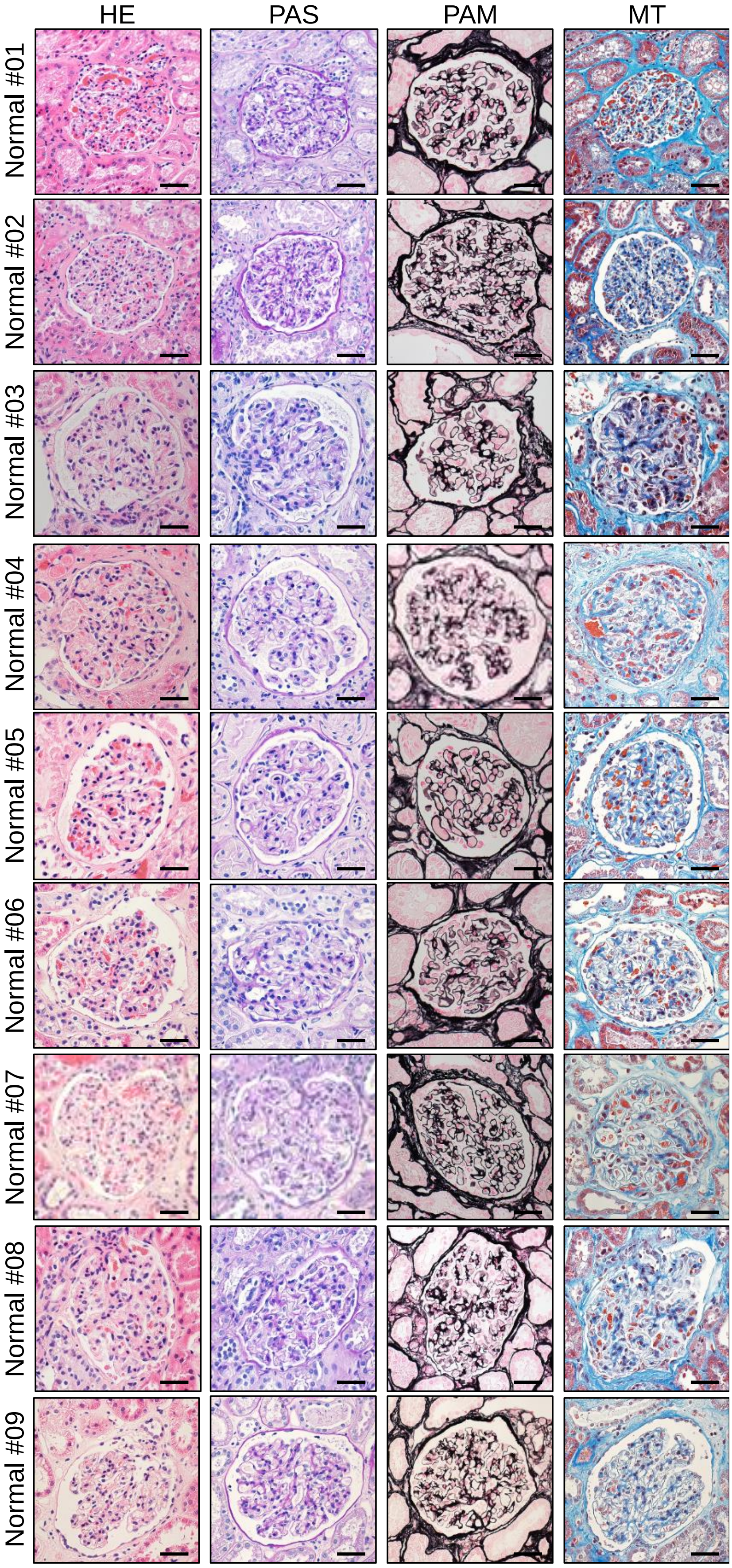
HE
PAS
PAM
MT
Normal #01
Normal #02
Normal #03
Normal #04
Normal #05
Normal #06
Normal #07
Normal #08
Normal #09

(b)

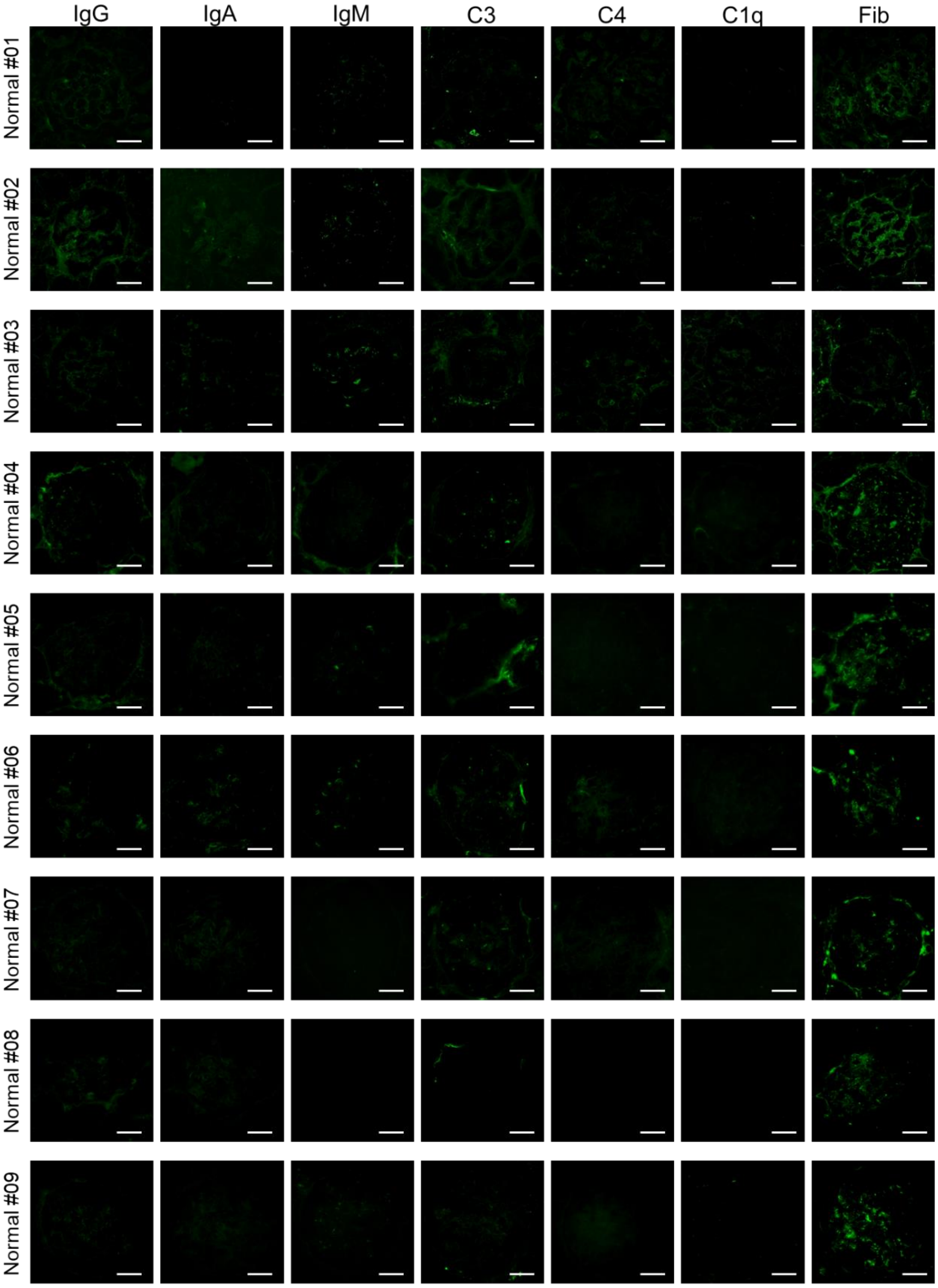
IgG
IgA
IgM
C3
C4
C1q
Fib
Normal #01
Normal #02
Normal #03
Normal #04
Normal #05
Normal #06
Normal #07
Normal #08
Normal #09

(c)

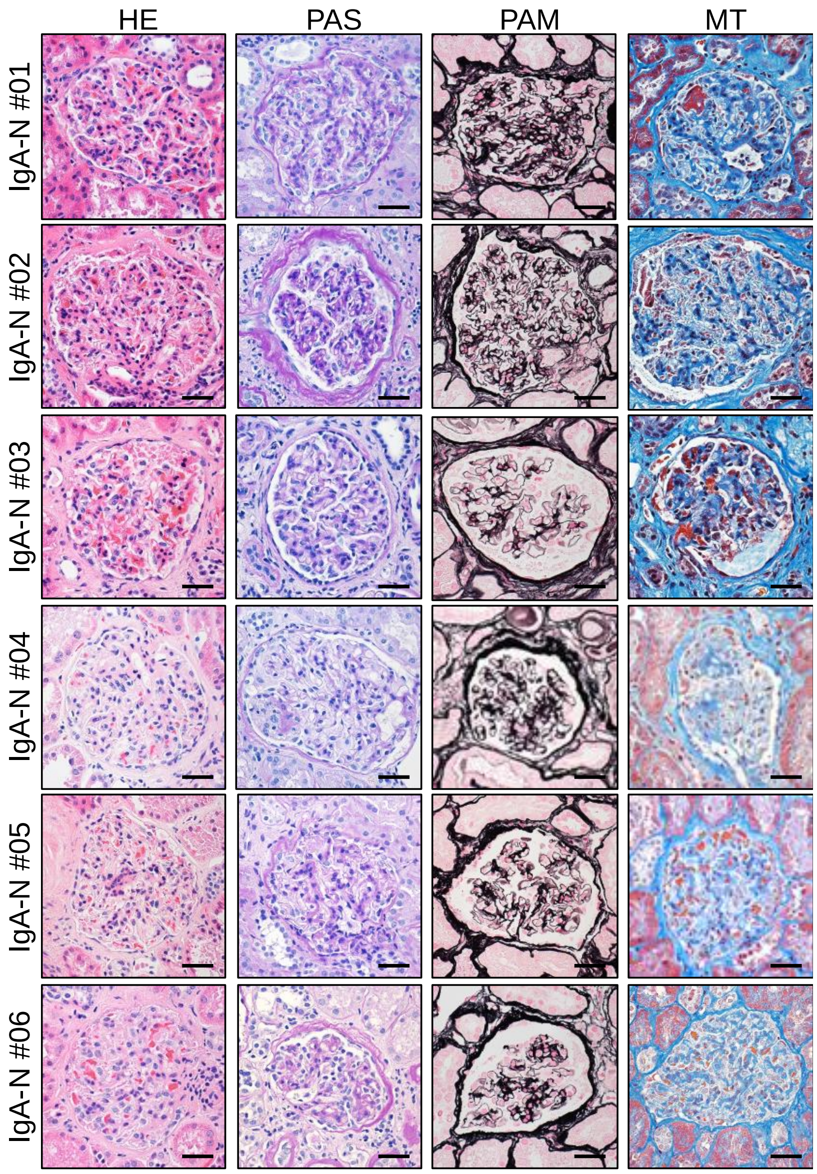

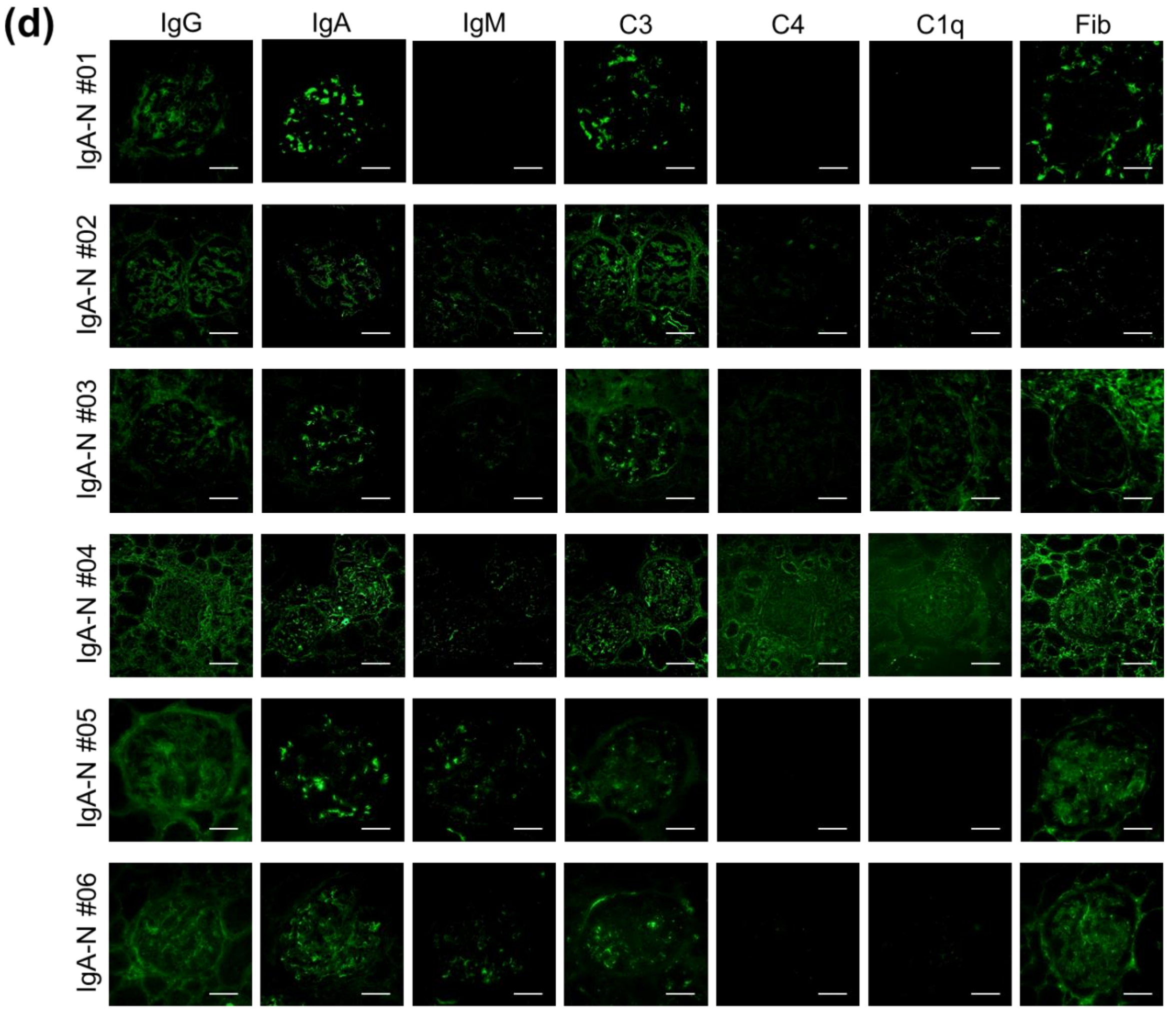


**Supplementary Figure 1: Histological staining images**

**(a)** Tissue staining image diagnosed as normal; **(b)** Immunofluorescence staining image of tissue diagnosed as normal; **(c)** Tissue staining image diagnosed as IgA-N; and **(d)** Immunofluorescence staining image of tissue diagnosed as IgA nephropathy. Scale bars = 40 μm. IgA-N (Immunoglobulin A nephropathy), HE (Hematoxylin& Eosin staining), PAS (periodic acid-Schiff staining), PAM (periodic acid-methenamine silver staining), MT (Masson's trichrome staining), IgG (Immunoglobulin G), IgA (Immunoglobulin A), IgM (Immunoglobulin M), C3 (Complement c3), C4 (Complement c4), C1q (Complement c1q), Fib (Fibrinogen).

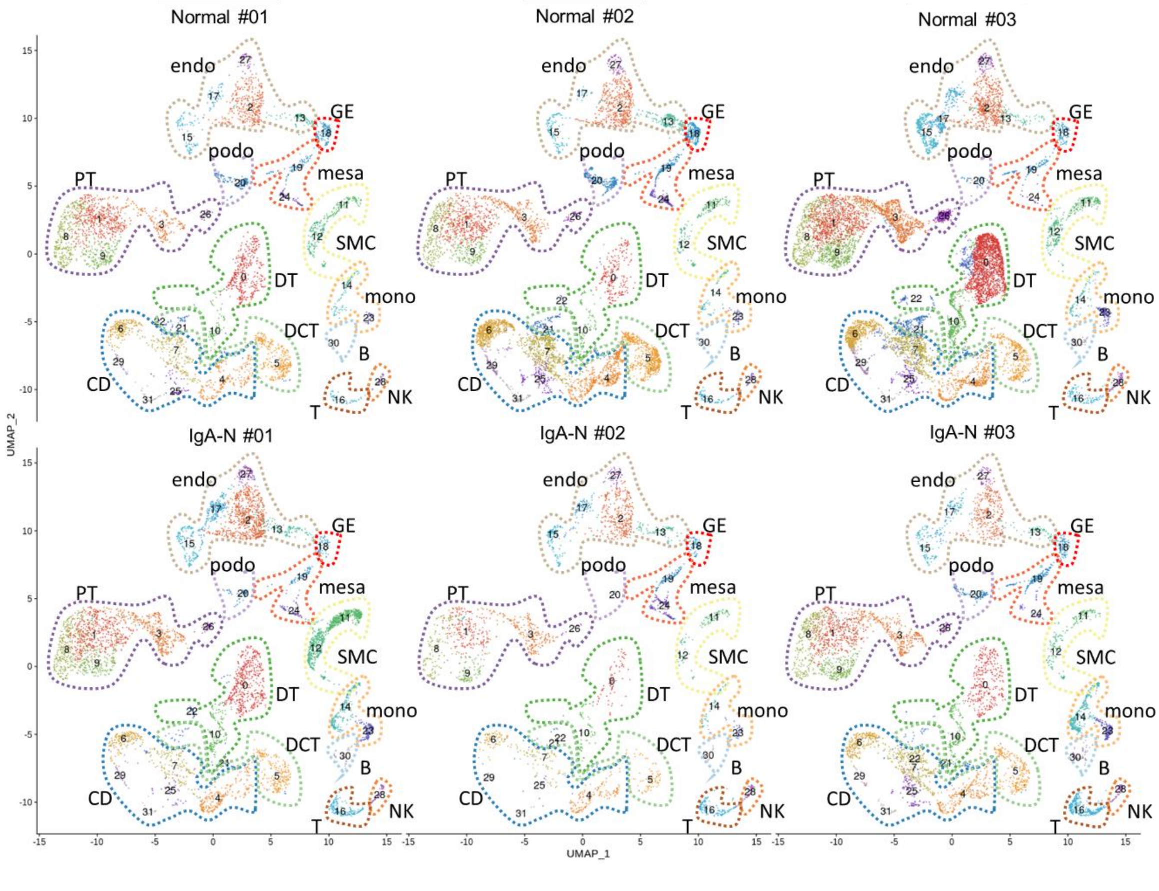


**Supplementary Figure 2: scRNA-seq data of cells isolated from each human renal tissue specimen. UMAP plots of major renal cell populations identified by unsupervised clustering and annotation with marker genes**

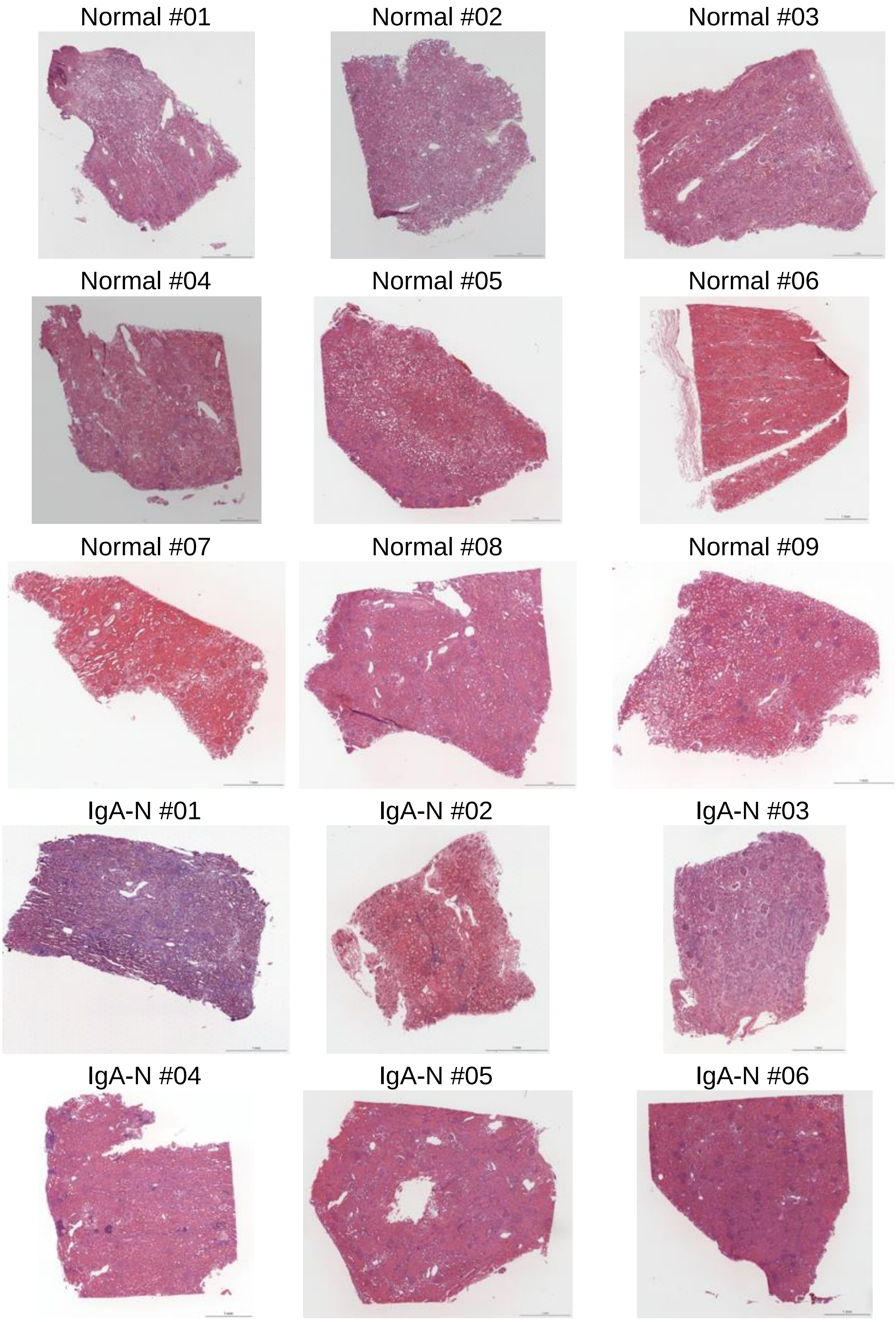


**Supplementary Figure 3: Visium HE images of each human renal tissue specimen**

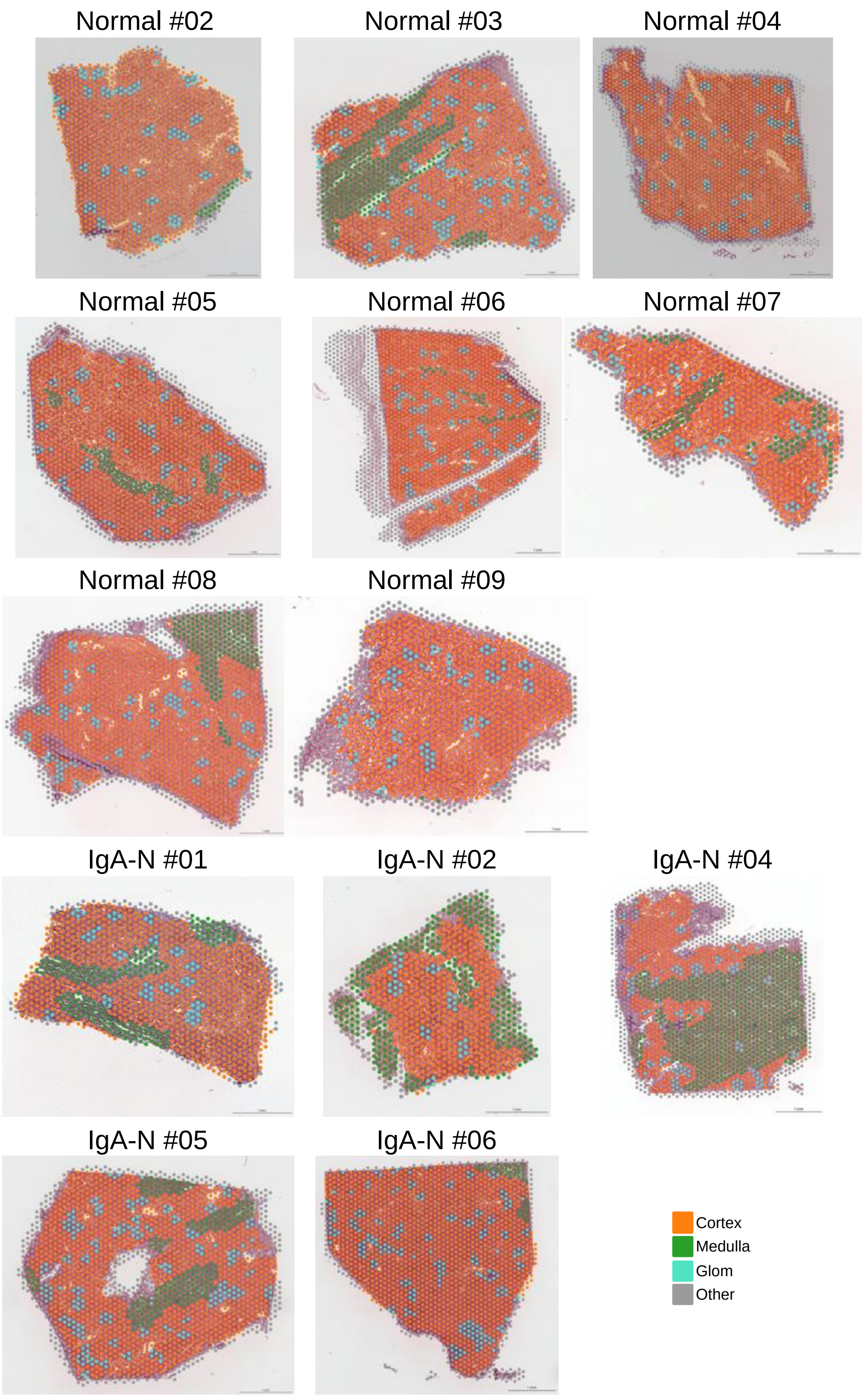


**Supplementary Figure 4: Visium data of spots on tissue from each human renal tissue specimen**

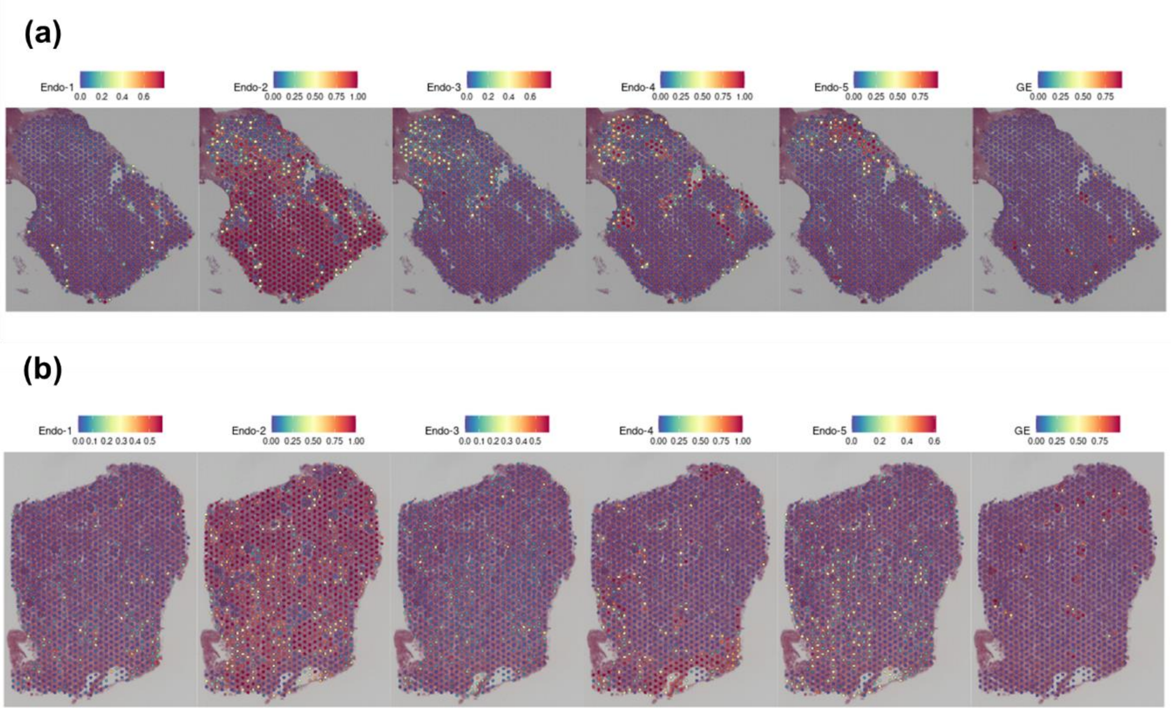


**Supplementary Figure 5: Integrated analysis of scRNA-seq and Visium.**

Predictive score for each spot on sections of **(a)** control specimens and **(b)** IgA-N specimens. Dot colors represent scaled predictive scores.

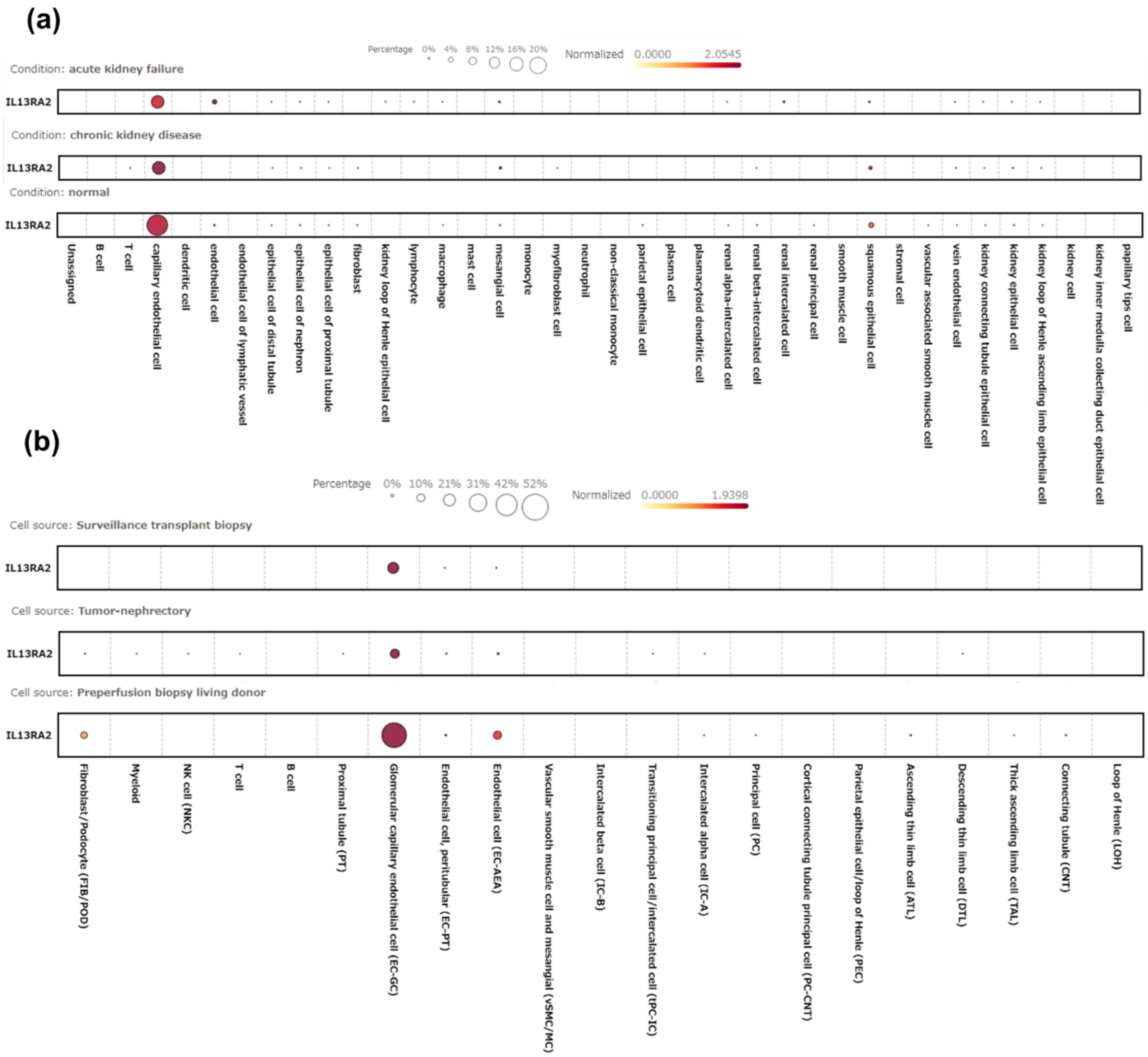
(a)
Percentage 0% 4% 8% 12% 16% 20%
Normalized 0.0000 2.0545
Condition: acute kidney failure
IL13RA2
Condition: chronic kidney disease
IL13RA2
Condition: normal
IL13RA2
Unassigned
B cell
T cell
capillary endothelial cell
dendritic cell
endothelial cell
endothelial cell of lymphatic vessel
epithelial cell of distal tubule
epithelial cell of nephron
epithelial cell of proximal tubule
fibroblast
kidney loop of Henle epithelial cell
lymphocyte
macrophage
mast cell
mesangial cell
monocyte
myofibroblast cell
neutrophil
non-classical monocyte
parietal epithelial cell
plasma cell
plasmacytoid dendritic cell
renal alpha-intercalated cell
renal beta-intercalated cell
renal intercalated cell
renal principal cell
smooth muscle cell
squamous epithelial cell
stromal cell
vascular associated smooth muscle cell
vein endothelial cell
kidney connecting tubule epithelial cell
kidney epithelial cell
kidney loop of Henle ascending limb epithelial cell
kidney cell
kidney inner medulla collecting duct epithelial cell
papillary tips cell
(b)
Percentage 0% 10% 21% 31% 42% 52%
Normalized 0.0000 1.9398
Cell source: Surveillance transplant biopsy
IL13RA2
Cell source: Tumor-nephrectory
IL13RA2
Cell source: Preperfusion biopsy living donor
IL13RA2
Fibroblast/Podocyte (FIB/POD)
Myeloid
NK cell (NKC)
T cell
B cell
Proximal tubule (PT)
Glomerular capillary endothelial cell (EC-GC)
Endothelial cell, peritubular (EC-PT)
Endothelial cell (EC-AEA)
Vascular smooth muscle cell and mesangial (vSMC/MC)
Intercalated beta cell (IC-B)
Transitioning principal cell/intercalated cell (tPC-IC)
Intercalated alpha cell (IC-A)
Principal cell (PC)
Cortical connecting tubule principal cell (PC-CNT)
Parietal epithelial cell/loop of Henle (PEC)
Ascending thin limb cell (ATL)
Descending thin limb cell (DTL)
Thick ascending limb cell (TAL)
Connecting tubule (CNT)
Loop of Henle (LOH)

(c)

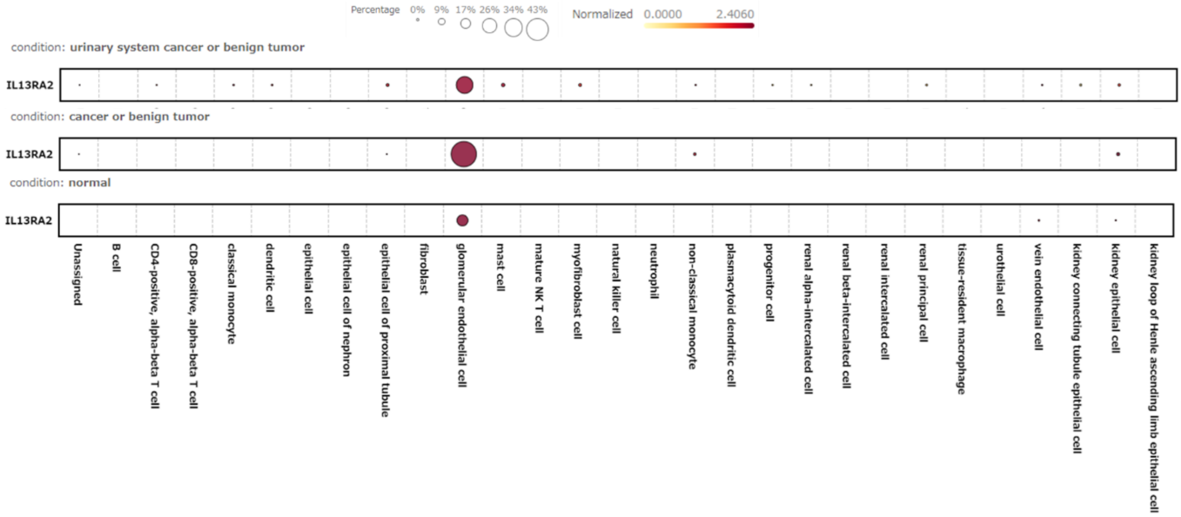


(d)

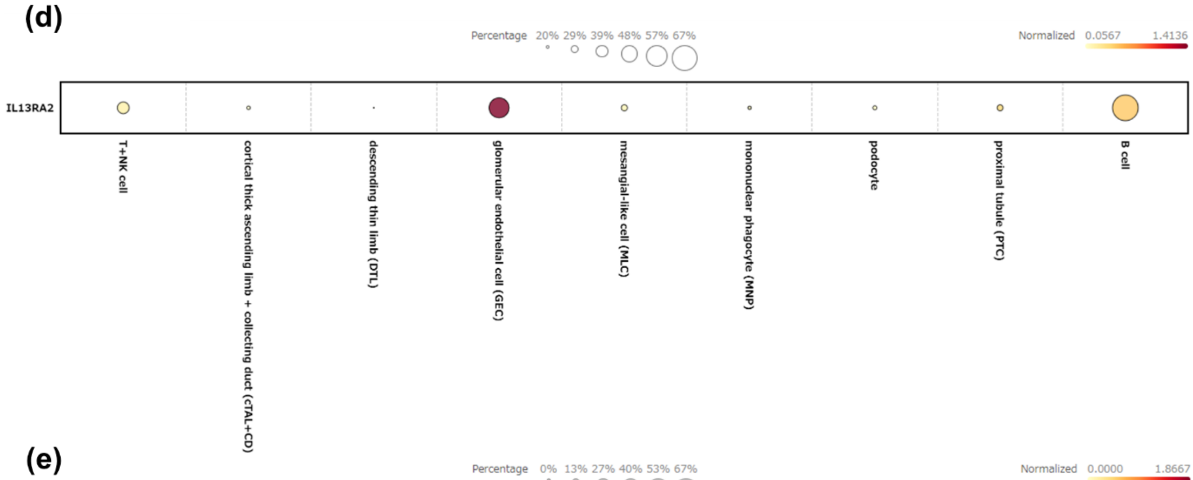


(e)

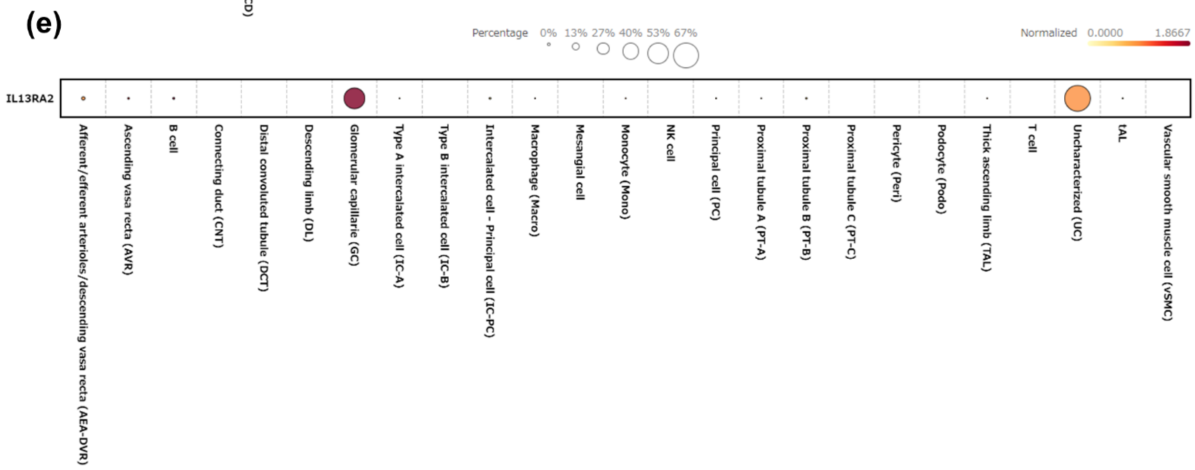


**Supplementary Figure 6: Confirmation of IL13RA2 expression in the publicly available data of human renal tissue scRNA-seq.**

All data show **(a)** LAKEETAL, **(b)** GSE140989, **(c)** PMID31604275_Mature, **(d)** GSE160048, and **(e)** GSE159115_6_BENIGN.

**Supplementary Table 1: Details of antibodies used in the study**

| Name | Source | Type |
|---|---|---|
| FITC IgG goat anti-human IgG FC | MP Biomedicals, #0855150 | Goat, polyclonal |
| FITC IgG goat anti-human IgA (α chain) | MP Biomedicals, #0855077 | Goat, polyclonal |
| Goat anti-human IgM-FITC | Bio-Rad, #STAR145F | Goat, polyclonal |
| FITC IgG goat anti-human complement C3 | MP Biomedicals, #0855167 | Goat, polyclonal |
| FITC IgG goat anti-human complement C4 | MP Biomedicals, #0855168 | Goat, polyclonal |
| FITC IgG goat anti-human complement C1q | MP Biomedicals, #0855166 | Goat, polyclonal |
| FITC IgG goat anti-human fibrinogen | MP Biomedicals, #0855169 | Goat, polyclonal |
| EHD3 polyclonal antibody | Proteintech, #25320-1-AP | Rabbit, polyclonal |
| Endothelin 1 monoclonal antibody (TR.ET.48.5) | Invitrogen, #MA3-005 | Mouse, monoclonal |
| Sclerostin polyclonal antibody | Proteintech, #21933-1-AP | Rabbit, polyclonal |
| IL-13RA2 polyclonal antibody | Proteintech, #11059-1-AP | Rabbit, polyclonal |
| Human CCL2/JE/MCP-1 antibody | R&D Systems, #MAB679-100 | Mouse, monoclonal |
| CXCL2 antibody | antibodies-online, #ABIN2473896 | Rabbit, polyclonal |
| EDNRB polyclonal antibody | Proteintech, #20964-1-AP | Rabbit, polyclonal |
| Goat anti-rabbit IgG(H+L) cross-adsorbed secondary antibody, Alexa Fluor™ 488 | Invitrogen, #A-11008 | Goat, polyclonal |
| Goat anti-rabbit IgG(H+L) | Invitrogen, #A-11011 | Goat, polyclonal |

| Name | Source | Type |
| --- | --- | --- |
| cross-adsorbed secondary antibody, Alexa Fluor™ 568 | | |
| Donkey anti-mouse IgG (H+L), Alexa Fluor™ 594 | abcam, #ab150108 | Donkey, polyclonal |

The other supplemental tables below are supplied by another file in csv format.

**Supplementary Table 2: General statistics of scRNA-seq data**

**Supplementary Table 3: Cluster-specific genes (scRNA-seq data)**

**Supplementary Table 4: General statistics of ST-seq data**

**Supplementary Table 5: DEGs for normal status versus pathological status for each cell type (scRNA-seq data)**

**Supplementary Table 6: Pathways in mesangial and GE**

**Supplementary Table 7: List of differentially expressed genes with a z-score of >2 contained in the inflammation-associated pathways in glomerular endothelial cells from IgA nephropathy specimens**

**Supplementary Table 8: DEGs in glomerular spots in IgA-N (ST-seq data)**

**Supplementary Table 9: Comparison analysis of mesangial and GE**
